# Genome-wide data for effective conservation of manta and devil ray species

**DOI:** 10.1101/458141

**Authors:** Jane Hosegood, Emily Humble, Rob Ogden, Mark de Bruyn, Si Creer, Guy Stevens, Mohammed Abudaya, Kim Bassos-Hull, Ramon Bonfil, Daniel Fernando, Andrew D. Foote, Helen Hipperson, Rima W. Jabado, Jennifer Kaden, Muhammad Moazzam, Lauren Peel, Stephen Pollett, Alessandro Ponzo, Marloes Poortvliet, Jehad Salah, Helen Senn, Joshua Stewart, Sabine Wintner, Gary Carvalho

## Abstract

Practical biodiversity conservation relies on delineation of biologically meaningful units, particularly with respect to global conventions and regulatory frameworks. Traditional approaches have typically relied on morphological observation, resulting in artificially broad delineations and non-optimal species units for conservation. More recently, species delimitation methods have been revolutionised with High-Throughput Sequencing approaches, allowing study of diversity within species radiations using genome-wide data. The highly mobile elasmobranchs, manta and devil rays (*Mobula* spp.), are threatened globally by targeted and bycatch fishing pressures resulting in recent protection under several global conventions. However, a lack of global data, morphological similarities, a succession of recent taxonomic changes and ineffectual traceability measures combine to impede development and implementation of a coherent and enforceable conservation strategy. Here, we generate genome-wide Single Nucleotide Polymorphism (SNP) data from among the most globally and taxonomically representative set of mobulid tissues. The resulting phylogeny and delimitation of species units represents the most comprehensive assessment of mobulid diversity with molecular data to date. We find a mismatch between current species classifications, and optimal species units for effective conservation. Specifically, we find robust evidence for an undescribed species of manta ray in the Gulf of Mexico and show that species recently synonymised are reproductively isolated. Further resolution is achieved at the population level, where cryptic diversity is detected in geographically distinct populations, and indicates potential for future traceability work determining regional location of catch. We estimate the optimal species tree and uncover substantial incomplete lineage sorting, where standing variation in extinct ancestral populations is identified as a driver of phylogenetic uncertainty, with further conservation implications. Our study provides a framework for molecular genetic species delimitation that is relevant to wide-ranging taxa of conservation concern, and highlights the potential for genomic data to support effective management, conservation and law enforcement strategies.

## Introduction

The Anthropocene has been characterised by unprecedented human exploitation of natural resources, resulting in global threats to biodiversity and extinction events across diverse taxa (Dirzo et al. 2014). Effective measures for biodiversity conservation require understanding and characterisation of diversity within and among species. The field of conservation genetics focuses on quantifying diversity across space and time (Allendorf et al. 2010), facilitated by increasingly powerful genome-wide data. Such genomic approaches also have applications in investigating the evolutionary processes generating biodiversity (Seehausen et al. 2014), providing further knowledge towards mitigating declines.

Biodiversity conservation is enacted through global conventions and regulatory frameworks implemented through legislation at the species level. Examples include the Convention on the International Trade in Endangered Species of Wild Fauna and Flora (CITES), and the Convention on the Conservation of Migratory Species of Wild Animals (CMS). In practice however, conservation initiatives and enforcement of regulations typically occur at a more local scale. Species therefore have two important impacts on conservation implementation; as units for inclusion in international conventions designed to coordinate conservation efforts, and representing identifiable targets against which conservation actions are directed and measured (Mace, 2004). Effective wildlife protection, management and law enforcement therefore depend on unambiguous classification of diversity into biologically relevant species units. Recent examples of proposed taxonomic revisions having far-reaching consequences for conservation include giraffe (Fennessy et al. 2016) and African elephant (Roca et al. 2001), where genetic research underpins possible reclassification and changes to the legal status of these megafauna.

Consequently, species delimitation, the process by which individuals are grouped into reproductively isolated and separately evolving units, is a fundamental application of genomic data to biodiversity conservation, with numerous methods available (Carstens et al. 2013; Grummer et al. 2014; Leache et al. 2014; Rannala 2015). Traditional approaches typically relied upon morphological observation, often resulting in artificially broad delineations arising from difficulties detecting and identifying cryptic species (Frankham et al. 2012), and impeding conservation efforts. More recently, DNA sequencing has allowed genetic data to be utilised for species delimitation, although interpretation may be challenging in recently diverged groups with substantial incomplete lineage sorting (Maddison & Knowles, 2006). Species delineations should minimise ambiguity by defining species units on the basis of reproductive isolation associated with limited gene flow and a lack of shared alleles (Frankham et al. 2012) and may therefore be optimised with evaluation of genomic data (Shafer et al. 2015). Genome-wide multi-locus approaches have increased the resolution of species delimitation studies, clarified contentious relationships and phylogenies (Leache et al. 2014; Herrera & Shank, 2016), disclosed previously unknown diversity (Pante et al. 2014) and elucidated evolutionary processes (Foote & Morin, 2016; Campbell et al. 2018). In addition, there are further applications in characterisation of Conservation Units and Evolutionary Significant Units to further enhance conservation efforts (Funk et al. 2012).

The importance of judiciously defined species or management units is particularly apparent in fisheries management (Reiss et al. 2009). Overexploitation of marine fisheries is a global problem (Agnew et al. 2009) resulting in loss of genetic diversity and bottlenecks in many species (Hauser et al. 2002; Pinsky & Palumbi, 2014). One group of heavily targeted fishes are the manta and devil rays (*Mobula* spp.; collectively, mobulids). Despite substantial economic value through tourism (O’Malley et al. 2013), these highly-mobile, circumglobally distributed megafauna are threatened by intense targeted and bycatch fishing pressure driven by demand for gill plates (Couturier et al, 2012; O’Malley et al. 2017). Consumptive exploitation of manta and devil rays is considered unsustainable due to slow life history traits, hindering recovery from fishing impacts (Dulvy et al. 2014; Croll et al. 2016). To alleviate threats, all mobulid species are listed on CITES Appendix II to regulate international trade, and on CMS Appendices I and II to coordinate protection and implement conservation efforts. These fish are poorly studied however, and marked homogeneity in morphology among species, a lack of representative global samples and population-level data, ongoing taxonomic debate, and ineffectual traceability measures constrain classification of optimal species units for conservation (Stewart et al. 2018). Understanding of evolutionary history and diversification in the Mobulidae derives from few studies, which indicate secondary contact and introgression among lineages may further impede efforts to delimit species boundaries (Kashiwagi et al. 2012; Poortvliet et al. 2015).

Recent evaluation of eleven previously recognised mobulid species across two genera recognised eight species, and called for the genus *Manta* (manta rays; *Manta alfredi* and *Manta birostris*) to be subsumed into *Mobula* (devil rays) (White et al. 2017). Other recent taxonomic changes include the resurrection of *Manta alfredi*; recognising two species of manta ray (Marshall et al. 2009; Kashiwagi et al. 2012), yet evidence remains of historic (Kashiwagi et al. 2012) and modern (Walter et al. 2014) hybridisation. In addition, a third putative species of manta ray is hypothesised to occur in the Caribbean (Marshall et al. 2009; Hinojosa-Alvarez et al. 2016). To date however, studies have relied on morphological observation (Notarbartolo Di Sciara 1987; Marshall et al. 2009; White et al. 2017) and/or been limited to evaluation of a handful of genetic markers, with heavy reliance on mitochondrial DNA (Kashiwagi et al. 2012; Hinojosa-Alvarez et al. 2016). Previous studies have also been geographically restricted and reliant on few samples (White et al. 2017), resulting in classifications that fail to encapsulate the extent of diversity within the group and compromise the effectiveness of conservation efforts.

Here, we generate double-digest Restriction-site Associated DNA sequence (ddRAD) data from the largest and most comprehensive set of mobulid tissue samples available. We demonstrate utility in delimiting informative species units for conservation, detecting cryptic diversity, and improving our understanding of associated evolutionary processes in a global radiation of socio-economically important marine megafauna.

## Methods

### Sampling and Sanger sequencing

Tissue samples were obtained from existing collections and sampling initiatives of researchers and organisations worldwide, yielding samples representing all mobulid species from a broad geographical range (Figure 1 and Supplementary Table 1), including *Mobula japanica, Mobula eregoodootenkee* and *Mobula rochebrunei,* currently considered junior synonyms of *Mobula mobular*, *Mobula kuhlii* and *Mobula hypostoma,* respectively (White et al. 2017), and an outgroup, *Rhinoptera bonasus.* Samples were identified to species based on characteristics described by Stevens et al. (2018), using original species names assigned and valid at the time of collection.

**Figure 1:**
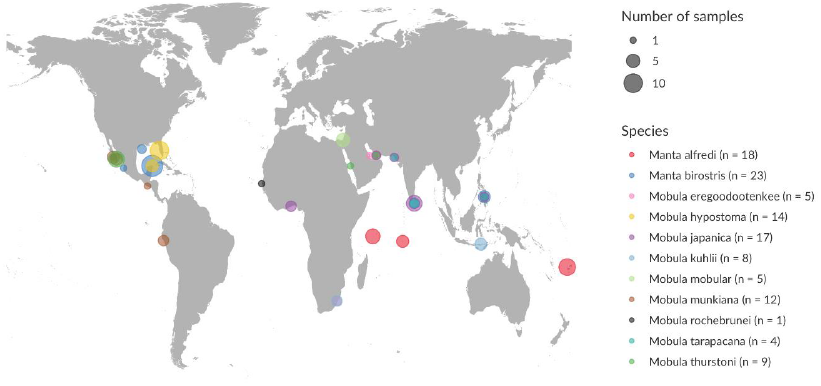
Sampling locations. Species are represented by coloured points, scaled for sample size. Total numbers of samples for each species provided in the key. Further details in Supplementary Table 1. Species names are those assigned at time of collection, some now considered invalid (White et al. 2017).

Genomic DNA was extracted using the Qiagen DNeasy Blood and Tissue Kit and DNA yield measured using a Qubit 3.0 Broad Range Assay. Extracts were quality assessed on 1% agarose gels stained with SafeView. The single sample of *Mobula rochebrunei* was from a museum specimen stored in formalin and yielded no detectable DNA.

To evaluate traditional markers for mobulid species delimitation, PCR amplification of an approximately 650bp portion of the Cytochrome Oxidase Subunit I (COI) gene was performed using universal Fish primers (Ward et al. 2005). Where these primers failed (for *M. munkiana* and *M. hypostoma*), primers MunkF1 (GGGATAGTGGGTACTGGCCT) and MunkR1 (AGGCGACTACGTGGGAGATT) were designed using Primer-BLAST (Ye et al. 2012). 15 µl PCR reactions consisted of: 5.6 µl nuclease-free water, 7.5 µl ReddyMix PCR Master Mix (ThermoFisher), 0.45 µl of each primer and 1 µl DNA. PCR cycling conditions were: 95°C for 2 min, 35 cycles of 94°C for 30s, 54°C for 30s and 72°C for 1 min and final extension of 72°C for 10 mins. Sanger sequencing was conducted by Macrogen Europe. Data was aligned using ClustalW and the alignment checked for stop codons in MEGA7 (Kumar et al. 2016). The HKY+G model was identified as optimal for our COI dataset using the Find Best Model option in MEGA7. A Maximum Likelihood tree was built with 1,000 bootstrap replicates.

### ddRAD library preparation and sequencing

ddRAD libraries were prepared using a modified version of the original protocol (Peterson et al. 2012; see Palaiokostas et al. 2015) with restriction enzymes *SbfI* and *SphI* (NEB). Unique P1 and P2 barcode combinations were ligated to resulting DNA fragments, which were then size-selected between 400-700bp using gel electrophoresis and PCR amplified. A pilot ddRAD library was sequenced on Illumina MiSeq at the Institute of Aquaculture, University of Stirling. Subsequent ddRAD libraries were sequenced by Edinburgh Genomics© on Illumina HiSeq High Output v4, 2 x 125PE read module.

### Data quality control and filtering

Data quality was assessed with FastQC (Andrews, 2010), and processed in Stacks version 1.46 (Catchen et al. 2011). The process_radtags.pl module in Stacks was used to demultiplex the data, filter for adaptor sequences (allowing two mismatches), remove low quality sequence reads (99% probability) and discard reads with any uncalled bases. To minimise linkage disequilibrium in the SNP data, only forward reads were retained for subsequent analyses. Short fragments not removed through size-selection were filtered with a custom bash script (8.5% of reads).

The denovomap.pl program in Stacks was used to assemble loci and call SNPs. The three main parameters for assembly were those generating the largest number of new polymorphic loci shared across 80% of individuals, following Paris et al. (2017). Four identical reads were required to build a stack (-m), stacks differing by up to four nucleotides were merged into putative loci (-M) and putative loci across individuals differing by up to five nucleotides were written to the catalog (-n), giving an average coverage of 105x. We then used the populations.pl program in Stacks to generate two VCF files containing SNPs present in at least 10 and 90 individuals, respectively. To remove paralogous loci and mitigate for allele dropout (Arnold et al. 2013; Gautier et al. 2013), loci sequenced at greater than twice or less than one-third the standard deviation of coverage, respectively, were identified and excluded using VCFtools (Danecek et al. 2011). The remaining loci were assessed for excess heterozygosity using VCFtools, and those exhibiting a significant probability of heterozygote excess were excluded. Finally, since Stacks ignores indels, SNPs in the last five nucleotide positions were assumed erroneous and excluded. The remaining loci and SNPs were written to a whitelist and filtered for a single random SNP per locus to minimise linkage using populations.pl. This resulted in two final SNP matrices, p10 and p90, with 7926 and 1762 SNPs and 47.1% and 14% missing data, respectively (Supplementary Table 2).

### Monophyly and clustering

Relationships among individuals were inferred through Maximum Likelihood phylogenetic analysis using RAxML version 8.2.11 (Stamatakis 2014). Both ddRAD datasets were analysed since missing data may influence aspects of phylogenetic inference (Leaché et al. 2015). The GTRGAMMA model of rate heterogeneity was implemented following assessment of best fit models in jModelTest (Darriba et al. 2015) and support assessed with 1,000 bootstrap replicates.

RAxML identified four highly supported clades separated by long branches. To assess how individuals cluster within these clades, dataset p10 was divided by clade (Supplementary Table 3) and Principal Components Analysis (PCA) performed on each using the R package Adegenet (Jombart 2008). After assessment of ten axes, three were retained in all cases. Populations.pl was used to calculate *F*ST values among inferred clusters.

### Bayes Factor Delimitation

Bayes Factor Delimitation (Leache et al. 2014) was conducted using the modified version of SNAPP (Bryant et al. 2012), implemented as a plug-in to BEAST version 2.4.8 (Bouckaert et al. 2014). The method allows for direct comparison of Marginal Likelihood Estimates (MLEs) for alternative species delimitation hypotheses, hereafter models, under the multispecies coalescent. Path sampling involved 10 steps (1,000,000 MCMC iterations, 20% burnin), implementing the log-likelihood correction. Since MLEs are affected by improper prior distributions, a gamma distribution was implemented on the lambda (tree height) parameter. To assess the effect of priors on the ranking order of models, models were also assessed retaining the default 1/X distribution on lambda, implementing upper and lower bounds (10,000 and 0.00001 respectively), for a proper prior. Bayes Factors (2logeBF) were calculated from the MLE for each model for comparison (Kass & Raftery 1995; Leache et al, 2014), as follows:

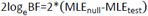

Positive 2logeBF values indicate support for the null model (<10 is decisive; Leache et al. 2014), negative values favour the tested model.

Due to computational constraints, dataset p90 underwent Bayes Factor Delimitation and the data were split by clade, as previously described, but including four random individuals from a sister species to evaluate support for interaction from higher phylogenetic levels. Alternative models were informed by the literature and analyses herein (Supplementary Tables 4-7). Models randomly assigning individuals to two or three species were assessed for each clade. Null models matched species defined by White et al. (2017).

### Species tree inference

Relationships among the Mobulidae were estimated through Maximum Likelihood phylogenetic analysis of both ddRAD datasets as above with RAxML (Stamatakis 2014). Consensus sequences for each species unit were ascertained using populations.pl in Stacks, providing a population map assigning individuals to optimal species units based on our previous analyses.

To test tree topology and evaluate uncertainty due to incomplete lineage sorting, species trees were additionally evaluated with SNAPP (Bryant et al. 2012), allowing each SNP to have its own history under the multispecies coalescent, whilst bypassing the need to sample individual gene trees. Due to the computational capacity required to run SNAPP, three individuals per species were randomly selected from dataset p90 whilst maximising geographical coverage within species. Random sampling of individuals with replacement was repeated a further three times, resulting in four subsampled alignments (Supplementary Table 8). MCMC chains consisted of 5,000,000 iterations, sampling every 1,000 and retaining default priors on lambda and theta for each independent analysis. Convergence to stationary distributions were observed after 20% burnin in TRACER (Rambaut et al. 2018), the distribution of trees visualised in DensiTree (version 2.2.6; Bouckaert 2010) and maximum clade credibility (MCC) trees drawn using TreeAnnotator (version 2.4.7; Bouckaert et al. 2014). Alternative prior combinations produced highly concordant results.

Multispecies coalescent based approaches assume that any discordance of topologies among loci results from incomplete lineage sorting, and do not consider introgression as a source of discordance. TreeMix (Pickrell & Pritchard, 2012) was applied to dataset p10 to evaluate evidence for significant introgression events within the Mobulidae by investigating the extent to which variation between user-defined groups is explained by a single bifurcating tree. Given uncertainty identified using SNAPP, specifically regarding the placement of *M. mobular*, the three-population test (Reich et al. 2009) was additionally used to test for ‘treeness’ between clades. Similar to TreeMix, the three-population test estimates covariance of allele frequencies among groups, but is simpler and less parameterised; potentially more powerful for identifying introgression. In addition to *M. mobular, M. alfredi* and *M. thurstoni* were selected randomly to represent their respective clades.

## Results

### Monophyly and clustering

Maximum Likelihood phylogenetic trees based on two genome-wide SNP matrices were highly congruent (Figure 2 and Supplementary Figure 1). Species groups formed well-supported clades separated by long branches. Principal Components Analyses (PCA) within each clade mirrored patterns in phylogenetic trees (Figure 3). Putative species, including recently synonymised species *Mobula kuhlii* and *Mobula eregoodootenkee* formed both reciprocally monophyletic groups with high bootstrap support (Figure 2) and tight clusters separated along axes explaining large portions of variance (63%-74%; Supplementary Figure 2). Two reciprocally monophyletic groups were detected within *Manta birostris*; an Atlantic and a global group, respectively (Figure 2), visible as clusters through PCA (Figure 3A). One individual was equally well, albeit poorly, placed with each clade in the two phylogenetic analyses (Fig 2 and Supplementary Figure 1) and in an intermediate position through PCA (Figure 3A). *Mobula japanica* and *Mobula mobular* formed a single monophyletic group with 100% bootstrap support (Figure 2), with no clear separation through PCA (Figure 3C-D). Whilst the first axis provides limited evidence to suggest a clustering of individuals into Indo-Pacific and Atlantic (including Mediterranean) groups, this explained only 8.6% of variance (Supplementary Figure 2E), with minimal differentiation between these two clusters (*F*ST = 0.061). Geographically separated populations of *Manta alfredi* and *Mobula kuhlii* formed highly-supported monophyletic groups (Figure 2) and were demarcated clearly through PCA (Figure 3B; Figure 3F), showing a high degree of differentiation (*F*ST = 0.16 and *F*ST = 0.32, respectively). COI sequences failed to achieve resolution sufficient to discriminate putative species, and phylogenetic analysis showed several multifurcating nodes (Supplementary Figure 3).

**Figure 2:**
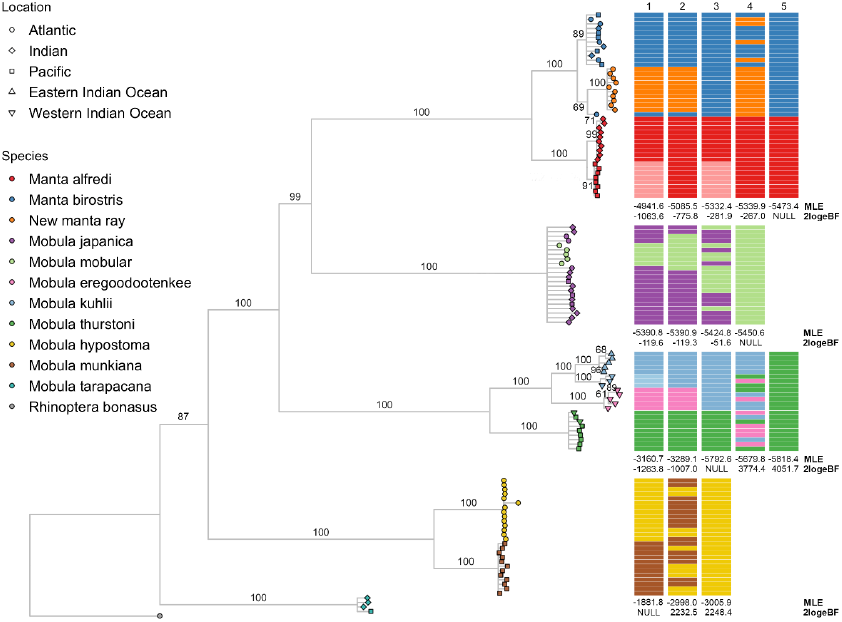
(Left) Maximum Likelihood Phylogenetic Tree of mobulid individuals based on 7926 SNPS (dataset p10). Coloured points indicate putative species, and shape indicates geographic origin of samples as specified in the key. Bootstrap values are shown on the branches and nodes with less than 50% support are collapsed. (Right) Bayes Factor Delimitation (BFD*) models with individuals assigned to species groups indicated by coloured bars are also presented, ranked in order of performance from left to right. Marginal Likelihood Estimates (MLEs) and Bayes Factors relative to the null model (2logeBF) are shown beneath each model for chains with a gamma prior on lambda. Models including individuals from a sister clade are not shown, as these consistently performed poorly. Species names are those assigned at time of collection, some now considered invalid (White et al. 2017).

**Figure 3:**
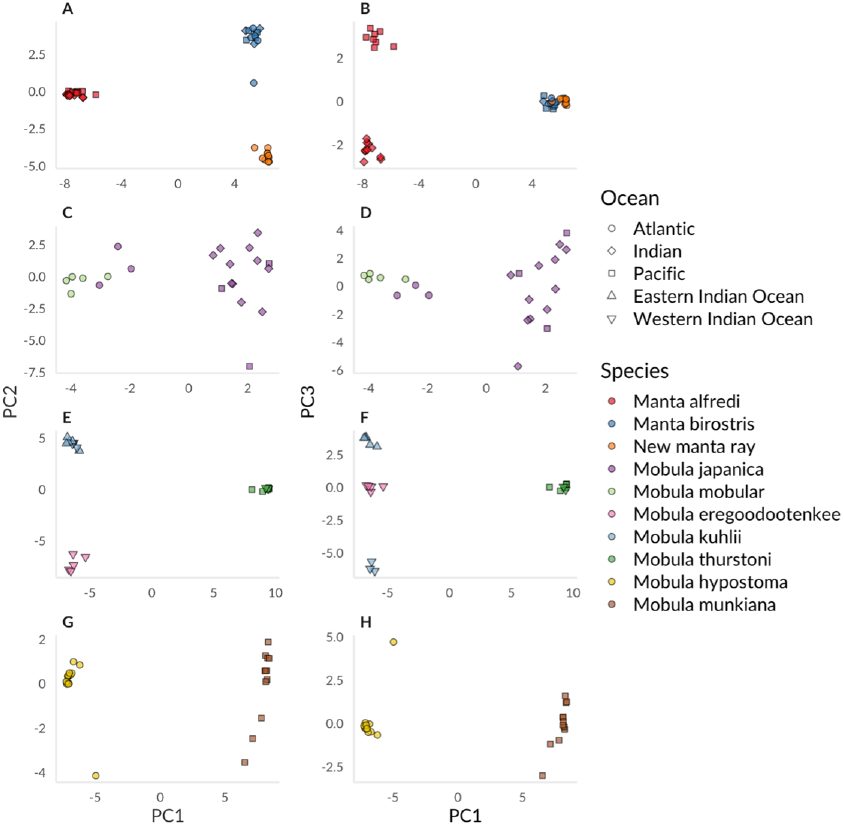
Principal Components 1-3 plotted for each mobulid clade. Individuals are represented by a point, colour indicates putative species, and shape indicates geographic origin of samples as specified in the key. Manta rays, A) PC1 and 2, and B) PC1 and 3; *M. mobular* and *M. japanica*, C) PC1 and 2, and D) PC1 and 3; *M. thurstoni*, *M. kuhlii* and *M. eregoodootenkee,* E) PC1 and 2, and F) PC1 and 3; *M. hypostoma* and *M. munkiana,* G) PC1 and 2, and H) PC1 and 3. Species names are those assigned at time of collection, some now considered invalid (White et al. 2017).

### Species Delimitation

Species models were compared within clades using Bayes Factor Delimitation (Figure 2). Marginal Likelihood estimates were unaffected by lambda priors, with no change in the rank order of models (Supplementary Tables 4-7). We find decisive support for models recognising the Gulf of Mexico and global *M. birostris* groups as separate species (2logeBF = −775.82; hereafter ‘*Mobula* sp. 1’ and ‘*M. birostris*’ respectively) and where individuals identified as *M. eregoodootenkee* belong to a separate species to *M. kuhlii* (2logeBF = −1007.04). Models splitting *M. mobular* and *M. japanica* based on geographic origin marginally out-performed the null model. Geographically informed models involving *M. alfredi* and *M. kuhlii* also performed well, achieving decisive support (2logeBF = −1063.58 and −1263.8, respectively). The null model was favoured within the *M. hypostoma* and *M. munkiana* clade. Models assessing support for interaction from higher levels and testing random individual assignments performed comparatively poorly (Supplementary Tables 4-7).

### Relationships among species

Maximum Likelihood species trees based on two genome-wide SNP matrices were highly congruent (Figure 4 and Supplementary Figure 4). Consistent with White et al. (2017), manta rays were nested within the genus *Mobula*, sister to *M. mobular* (≥98% bootstrap support) and hereafter all species of manta ray are referred to as *Mobula*. These trees strongly suggest that an undescribed third species of manta ray is sister to *M. birostris* (100% bootstrap support). *M. tarapacana* was tentatively placed on the group’s oldest lineage (84% bootstrap support).

**Figure 4:**
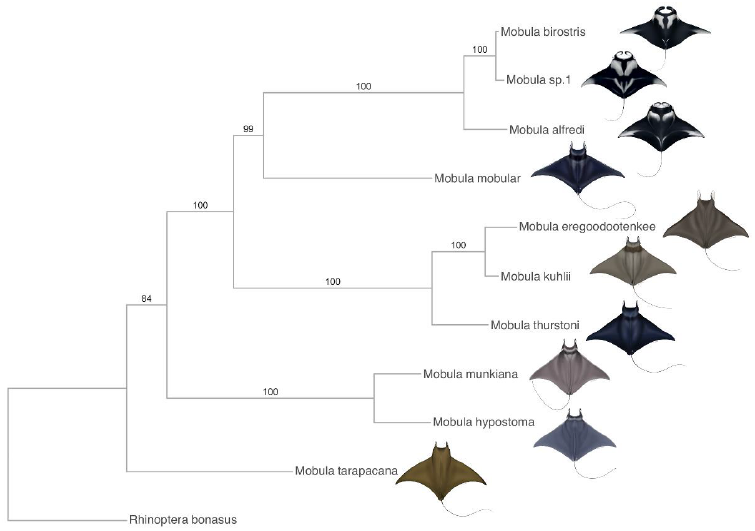
Maximum Likelihood tree of inferred mobulid species units based on 7902 SNPs (dataset p10). Bootstrap values are shown on the branches. The drawing of *Mobula* sp. 1 is based on images of dozens of individuals off the Yucatan Peninsula, Gulf of Mexico. Illustrations © Marc Dando.

Consensus species trees estimated under the multispecies coalescent exhibited relatively consistent topologies and theta estimates across independent runs, suggesting no major effect of subsampling on species tree topology inferred with SNAPP. *M. tarapacana* was consistently sister to *M. hypostoma* and *M. munkiana* (highest posterior density (HPD) = 1.0). Topological uncertainty at other nodes is apparent with a cloudogram of gene trees sampled from the posterior distribution (Figure 5 and Supplementary Figures 5-7). Relationships between sister species within clades remained consistent in alternative topologies within the 95% HPD, but large discrepancies in the placement of *M. mobular* (including *M. japanica*) relative to other clades were observed (Supplementary Table 9).

**Figure 5:**
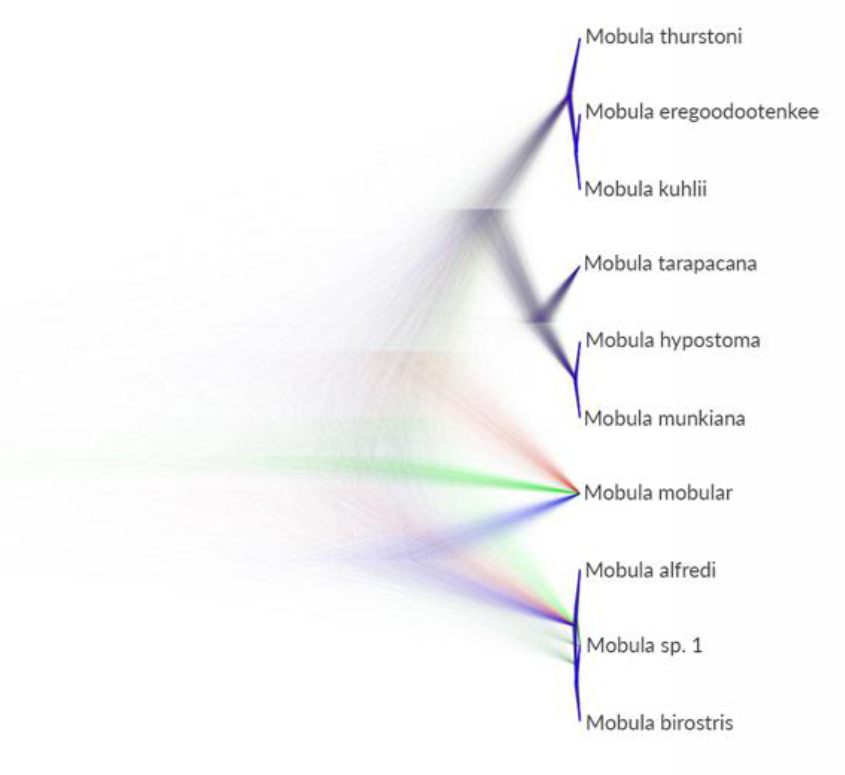
SNP phylogeny of 30 individuals assigned to ten species units based on 1242 SNPs (dataset p90, individual subsample 1; Supplementary Table 8). Tree cloud of sampled trees produced using DENSITREE (representing samples taken every 1000 MCMC steps from 5,000,0000 iterations) from SNAPP analysis to visualise the range of alternative topologies.

TreeMix inferred an admixture graph similar to trees produced with RAxML (Supplementary Figure 8), explaining 99.86% of variance, indicating mobulid species placement is unaffected by admixture. We found no evidence of introgression between clades containing *M. alfredi, M. mobular* and *M. thurstoni,* through three-population tests (Supplementary Table 10).

## Discussion

Genome-wide SNP data provide unprecedented resolution in a group of conservation concern, and our analyses produced the most extensive phylogeny for the Mobulidae to date. In contrast to previous studies examining mobulid diversity, the global nature of our dataset allowed us to identify reproductive isolation between lineages and distinguish between population and species units (Sukumaran & Knowles, 2017). We find a mismatch between current classifications and species units optimal for conservation, with implications for management and law enforcement. We provide robust evidence for a new species of manta ray and demonstrate that individuals identified as recently synonymised species *Mobula kuhlii* and *Mobula eregoodootenkee* are distinct and reproductively isolated. We therefore recommend that such units coincide with enforceable protection (see Appendix 1 for critical evaluation). In addition, we detect cryptic diversity between geographically segregated populations of *Mobula alfredi* and *Mobula kuhlii*, which may merit independent management.

These findings have international implications for practical conservation of the Mobulidae since legislation applies to species units and can severely impact anthropogenic pressures on wildlife populations. Our data suggest that the oceanic manta ray (*M. birostris*) and an undescribed species of manta ray (*Mobula* sp. 1) occur in sympatry in the Gulf of Mexico, since samples collected within sites fall into both groups, and provides evidence of hybridisation between these species (Figure 2; Fig 3A). Management of these similar species as independent units will therefore be challenging, potentially requiring blanket protection of all manta rays in regions where sympatry and/or hybridisation occur, and indeed such protection already exists in Mexico. Notwithstanding, *Mobula* sp. 1 is likely to occur over a broad geographic range, given patterns of distribution of its closest relatives. To establish effective conservation and traceability measures for this new species, it will therefore be necessary to formally describe *Mobula* sp. 1 and determine the extent of its range, which may extend into international waters or span areas with high fishing pressure lacking suitable protective measures.

Similarly, *Mobula eregoodootenkee* (as formerly recognised), shown here to be distinct from *M. kuhlii*, shares a geographic range with the latter across a region with intense fishing pressure (Notarbartolo di Sciara et al. 2017). Inference from related species suggests low reproductive output likely resulting in population sizes vulnerable to exploitation (Dulvy et al. 2014; Croll et al. 2016). It is therefore imperative that such units are managed separately. In contrast, species such as *M. mobular* may be of lower conservation priority given that *M. japanica* is a junior synonym (White et al. 2017; this study - see Appendix 1). Significant population structure in *M. alfredi* and *M. kuhlii* indicates potential for future traceability work to determine regional location of catch in these species (Appendix 1), which is increasingly required to comply with global obligations (Nielsen et al. 2012). Additional population-level studies will allow further assessment of stock structure within fisheries and delineation of mobulid conservation units for effective management.

We find substantial uncertainty in the placement of *M. mobular,* and trees within the 95% HPD where the manta rays (formerly genus *Manta*) are nested within *Mobula* are present in approximately equal proportions to trees where the former genera are reciprocally monophyletic (Supplementary Table 9). In groups that have undergone rapid speciation with large ancestral effective population size, the effects of incomplete lineage sorting on species tree estimation are particularly prominent (Lischer et al. 2014; Flouri et al. 2018). The Mobulidae have undergone recent rapid bursts of speciation (Poortvliet et al. 2015), and our estimates of mutation-scaled effective population size were larger on deeper branches of the tree, indicating large effective population size of extinct shared ancestral species (Supplementary Figure 9). Thus, standing variation in ancestral populations of mobulid rays is likely to drive uncertainty with respect to the validity of the genus *Manta*. Since we find no evidence of admixture driving these patterns, this uncertainty can be attributed to incomplete lineage sorting. Factors such as similarities in life history and difficulties distinguishing between related species in trade can lead to whole genera being listed on international conventions such as CITES designed to preserve biodiversity. Our data therefore demonstrates the importance of understanding the extent and nature of incomplete lineage sorting for effective conservation of threatened groups.

Genomic approaches are increasingly informative for inferring phylogenetic relationships among species. Results must, however, be interpreted with caution. Our Maximum Likelihood analysis identified *M. tarapacana* as the oldest mobulid lineage, coincident with similar analyses of nuclear data (White et al. 2017), yet our Bayesian analyses consistently placed *M. tarapacana* sister to *M. hypostoma* and *M. munkiana*; a previously unreported phylogenetic placement. Analyses employing mitochondrial data support *M. tarapacana* as sister to the manta rays and *M. mobular* (Poortvliet et al. 2015; White et al. 2017), an observation we were unable to reproduce. Discordant trees in phylogenomic studies may be attributed to few loci driven by positive selection resulting in convergent evolution, or evolutionary processes such as incomplete lineage sorting (Shen et al. 2017). Coalescent-based approaches, as applied here, account for the independent history of each gene tree and are therefore less likely to be influenced by single genes, highlighting the suitability of genome-wide data for the inference of species relationships.

Here, genome-wide data considerably enhances delimitation of species units for the conservation of manta and devil rays. These findings have profound implications for the practical conservation of a group threatened by fishing, and are relevant to enforcement of CITES regulations by laying the groundwork for species and regional traceability of parts in trade. Furthermore, we demonstrate the ability of genomic data to resolve and identify diversity within organismal radiations and improve understanding of evolutionary processes generating biodiversity. As such, this study provides a framework for molecular genetic species delimitation which is relevant to other wide-ranging taxa of conservation concern, and highlights the potential for applied research in supporting conservation, management and law enforcement.

## Acknowledgements

We are very grateful to the Save Our Seas Foundation (SOSF) and to The People’s Trust for Endangered Species (PTES) for providing generous support for this work. JH is supported by a NERC CASE studentship through the ENVISION DTP (CASE partner - Royal Zoological Society of Scotland) and has received additional grants from the Fisheries Society of the British Isles (FSBI) and the Genetics Society. Data analysis was supported by the UK Natural Environment Research Council (NERC) Biomolecular Analysis Facility at the University of Sheffield.

The authors are very grateful to the following people and organisations for their help and support sourcing and collecting tissue samples; J. Spaet, A. Moore, R. Brittain, G. Phillips, J. Schleyer, F. Doumbouya, D. Bowling, H. Pacey, BD. Croll, K. Newton, H. Badar Osmany, S. Hinojosa, all LAMAVE staff and volunteers, field team Captains D. Dougherty, P. Hull, G. Byrd, K. Wilkinson, B. DeGroot and organisations Akazul, West Africa Musee de la mer a Dakar, the Barefoot Collection and Planeta Oceano. We would also like to thank all the staff at Atlantis-The Palm Dubai for giving access to specimens brought in by fishermen and for their valuable help with data collection and dissections.

Blue Resources Trust (BRT) would like to thank the Department of Wildlife Conservation and the Department of Fisheries and Aquatic Resources for support provided to the fieldwork carried out in Sri Lanka. BRT also acknowledges the generous support provided by the SOSF and the Marine Conservation and Action Fund (MCAF) that enabled fieldwork in Sri Lanka.

We thank Disney Conservation Fund, SOSF and Mote Scientific Foundation for supporting sample collection in Florida. Special thanks also to the Local Government Unit of Jagna, the Philippines Bureau of Fisheries and Aquatic Resources Region 7. The SOSF D’Arros Research Centre is a main affiliate of the Seychelles Manta Ray Project, funded by the SOSF. Sample collection in the Seychelles was approved by, and conducted with the knowledge of, the Ministry of Environment, Energy, and Climate Change.

The National Commission for Fisheries and Aquaculture of Mexico (CONAPESCA) allowed RB the collection of samples in Mexico through research permit PPF/DGOPA-091/15; the National Commission for Natural Protected Areas (CONANP) of Mexico and authorities of the Biosphere Reserve of Whale Sharks kindly gave permission for work in the reserve. The SOSF and the MCAF provided funding for research in Mexico. The Perfect World Foundation generously funded RB for the replacement of a drone used to locate manta rays. The Mexican CITES authority, Secretary of Environment and Natural Resources (SEMARNAT) provided CITES export permit for tissue samples through permit MX 80544.

We also thank J. Taggart for his support with the ddRAD library preparation protocol, and for his help sequencing a pilot ddRAD library. G. Colucci assisted with DNA extractions and COI amplifications. In addition, we thank M. Dando for kindly agreeing for us to reproduce his illustrations.

AF was funded by the Welsh Government and Higher Education Funding Council for Wales through the Sêr Cymru National Research Network for Low Carbon, Energy and Environment, and from the European Union’s Horizon 2020 research and innovation programme under the Marie Skłodowska-Curie grant agreement No. 663830.

## Author Contributions

JH, EH, GC, MdB, RO, SC and GS designed and conceived of the study and secured funding for consumables relating to laboratory work. EH, GS, DF, AP, MA, JS, SP, SW, RJ, MP, MM, KBH, RB, JS and LP were responsible for sourcing and collecting samples. JH, HS and JK carried out laboratory work. JH, EH, GC, MdB, RO, SC, HH, AF and HS contributed to analysis of genome-wide SNP data. Figures were designed by EH and JH and produced by EH. All authors contributed to writing and editing the manuscript.

