## Supplementary Figures, Tables and Appendix for "Genome-wide data for effective conservation of manta and devil ray species"

### Supporting Information

**Supplementary Table 1**: Sample information. We use species names assigned to samples at the time of collection, some now considered invalid (White et al. 2017).

| Sample codes | Species | Location | Total | No. in COI Dataset | No. in ddRAD Datasets |
|---|---|---|---|---|---|
| 0130, 0131, 0132, 0135, 0136, 0138 | *Manta alfredi* | D'Arros, Amirante Islands, Seychelles | 6 | 6 | 6 |
| 0140, 0141, 0144, 0145, 0146, 0147, 0148, 0149 | *Manta alfredi* | Barefoot Channel, Yasawas Islands, Fiji | 8 | 8 | 8 |
| 0685, 0686 | *Manta alfredi* | Egmont, Chagos | 2 | 2 | 2 |
| 0687, 0688 | *Manta alfredi* | Diego Garcia, Chagos | 2 | 2 | 2 |
| 0731 | *Manta birostris* | Mirissa, Sri Lanka | 1 | 1 | 1 |
| 0732, 0736 | *Manta birostris* | Negombo, Sri Lanka | 2 | 1 | 2 |
| 0980[a], 0981[a], 0982[a], 0983[a], 0984[a], 0985, 0986, 0987, 0988, 1239[a], 1240[a], 1241[a], 1242[a] | *Manta birostris* | Yucatan Northern tip, Mexico Caribbean | 13 | 9 | 13 |
| 1102, 1110, 1114, 1122 | *Manta birostris* | Jagna, Bohol landing site, Philippines | 4 | 4 | 4 |
| 1168 | *Manta birostris* | Bahia de Banderas, Jalisco, Mexico Pacific | 1 | 1 | 1 |
| 1327[a], 1328[b] | *Manta birostris* | Flower Garden Banks National Marine Sanctuary, Texas, USA | 2 | 2 | 2 |

| | | | | | |
|---|---|---|---|---|---|
| 0684 | *Mobula eregoodootenkee* | Al Khor, Qatar | 1 | 1 | 1 |
| 0696 | *Mobula eregoodootenkee* | Arabian Gulf, United Arab Emirates | 1 | 1 | 1 |
| 0697 | *Mobula eregoodootenkee* | Fujeirah, United Arab Emirates | 1 | 1 | 1 |
| 0810, 0813 | *Mobula eregoodootenkee* | Zinkwazi, South Africa | 2 | 2 | 2 |
| 0873, 0874, 0886, 0888, 0891, 0900, 0924, 0929, 0933, 0938 | *Mobula hypostoma* | Sarasota, Florida | 10 | 10 | 10 |
| 0990, 0991, 0992, 0993 | *Mobula hypostoma* | Yucatan Northern tip, Mexico Caribbean | 4 | 4 | 4 |
| 0003, 0024 | *Mobula japanica* | Jagna, Bohol landing site, Philippines | 2 | 2 | 2 |
| 0219, 0229, 0234, 0283 | *Mobula japanica* | Negombo, Sri Lanka | 4 | 4 | 4 |
| 0300, 0329, 0343 | *Mobula japanica* | Mirissa, Sri Lanka | 3 | 3 | 3 |
| 0707, 0711 | *Mobula japanica* | Ras Al Khaimah, United Arab Emirates | 2 | 2 | 2 |
| 0757, 0771, 0773 | *Mobula japanica* | Lome, Togo | 3 | 3 | 3 |
| 0793 | *Mobula japanica* | Puerto Adolfo Lopez Mateos, Mexico Pacific | 1 | 1 | 1 |
| 0853, 0862 | *Mobula japanica* | Karachi, Pakistan | 2 | 2 | 2 |
| 0524 | *Mobula kuhlii* | Negombo, Sri Lanka | 1 | 1 | 1 |
| 0582, 0677 | *Mobula kuhlii* | Durban, South Africa | 2 | 2 | 2 |
| 0605 | *Mobula kuhlii* | Warner Beach, South Africa | 1 | 1 | 1 |
| 0774, 0776, 0777, 0778 | *Mobula kuhlii* | Maumere, Indonesia | 4 | 4 | 4 |
| 0101, 0104 | *Mobula mobular* | Gaza City Fishing Landing Site, Gaza, Palestine | 2 | 2 | 2 |

| | | | | | |
|---|---|---|---|---|---|
| 0111, 0112 | *Mobula mobular* | Khanyounis Fishing Landing Site, Gaza, Palestine | 2 | 2 | 2 |
| 0129 | *Mobula mobular* | Middle area fishing landing site, Gaza, Palestine | 1 | 1 | 1 |
| 0816, 0821, 0824, 0827 | *Mobula munkiana* | El Pardito, Mexico Pacific | 4 | 4 | 4 |
| 0830, 0833, 0840, 0843 | *Mobula munkiana* | Puerto Adolfo Lopez Mateos, Mexico Pacific | 4 | 4 | 4 |
| 0844 | *Mobula munkiana* | Buena Vista Port of San Jose, Guatemala | 1 | 1 | 1 |
| 1006, 1009, 1019 | *Mobula munkiana* | Peru | 3 | 3 | 3 |
| 0943 | *Mobula rochebrunei* | Musee de la Mer, Goree, Senegal | 1 | 0 | 0 |
| 0086 | *Mobula tarapacana* | Jagna, Bohol landing site, Philippines | 1 | 1 | 1 |
| 0408, 0411 | *Mobula tarapacana* | Mirissa, Sri Lanka | 2 | 2 | 2 |
| 0847 | *Mobula tarapacana* | Karachi, Pakistan | 1 | 1 | 1 |
| 0159 | *Mobula thurstoni* | Jeddah Fish Market, Saudi Arabia | 1 | 1 | 1 |
| 0712 | *Mobula thurstoni* | Fujeirah, United Arab Emirates | 1 | 1 | 1 |
| 0780, 0782, 0783, 0784, 0785, 0786, 0787 | *Mobula thurstoni* | El Pardito, Mexico Pacific | 7 | 7 | 7 |
| 1366, 1367, 1368, 1369, 1370 | *Rhinoptera bonasus* | Sarasota, Florida, USA | 5 | 0 | 5 |
| Totals | | | 121 | 110 | 120 |

[a] Samples identified as belonging to *Mobula* sp. 1 in our analyses
[b] Likely hybrid (*Manta birostris* x *Mobula sp. 1*)

**Supplementary Table 2**: Information on two SNP matrices generated using ddRAD, p10 and p90. The number of SNPs in individual level matrices are given, with numbers of SNPs in species level matrices in brackets.

| Dataset | Minimum individuals possessing a locus | Loci with >2x SD coverage | Loci with <1/3x SD coverage | Loci with >95% probability of heterozygote excess | SNPs | Missing Data |
|---|---|---|---|---|---|---|
| p10 | 10 | 1761 | 7661 | 24 | 7926 (7902) | 47% |
| p90 | 90 | 789 | 0 | 0 | 1762 (1755) | 14% |

**Supplementary Table 3**: Details of SNP matrices used to run PCAs. Dataset p10 (7926 SNPs) was split into clades for ease of visualisation, resulting in sites unsampled within clades to drop out.

| Clade | SNPs retained |
|---|---|
| *M. alfredi* and *M. birostris* (including *Mobula* sp. 1) | 5730 |
| *M. mobular* (including samples identified as *M. japanica*) | 5428 |
| *M. thurstoni* and *M. kuhlii* (including samples identified as *M. eregoodootenkee*) | 5384 |
| *M. hypostoma* and *M. munkiana* | 5063 |

**Supplementary Table 4**: Details of species delimitation models tested with Bayes Factor Delimitation (BFD*) within the first clade identified, the manta rays; *Manta alfredi* and *Manta birostris*, including individuals from sister clade *Mobula japanica* 0707, 0283, 0771 and 0024.

| Rank | SNPs retained | MLE ($2log_eBF$) - gamma prior | MLE ($2log_eBF$) - default 1/X prior | Description | Rationale | Reference |
|---|---|---|---|---|---|---|
| 1 | 1746 | -4941.58 (-1063.58) | -4946.44 (1054.72) | Split *M. alfredi* into two species units in Indian and Pacific Oceans. In addition, another species of manta ray is present in the Atlantic Ocean, sharing most recent common ancestor with *M. birostris.* | Monophyly of *M. alfredi* individuals in these ocean basins, and distinguishability through PCA. *M. birostris* split hypothesised in several studies. | This study |
| 2 | 1749 | -5085.46 (-775.82) | -5086.94 (-773.72) | In addition to *M. alfredi* and *M. birostris*, a third species of manta ray is present in the Atlantic Ocean, sharing most recent common ancestor with *M. birostris.* | Hypothesised in several studies. Some showing divergence on phylogenetic trees following this pattern | Marshall et al. 2009; Hinojosa-Alvarez et al. 2016; this study |
| 3 | 1748 | -5332.41 (-281.92) | -5335.11 (-277.38) | Split *M. alfredi* into two species units in Indian and Pacific Oceans, but recognise a single species within the *M. birostris* complex | Monophyly of *M. alfredi* individuals based on these ocean basins, and distinguishability through PCA. Potential hybridisation between *M. birostris* groups. | This study |

| | | | | | | |
|---|---|---|---|---|---|---|
| 4 | 1750 | -5339.87 (-267) | -5341.8 (-264) | Split *M. birostris* based entirely on geography (Atlantic/Gulf of Mexico individuals a separate species unit). Single species within *M. alfredi.* | To assess whether all *M. birostris* individuals sampled in the Atlantic could be considered a distinct species from *M. birostris* sampled elsewhere | Marshall et al. 2009 |
| 5 | 1751 | -5473.37 (Null) | -5473.8 (Null) | Null model and current arrangement – two species of manta ray: *M. alfredi* and *M. birostris* | Current taxonomic arrangement | Marshall et al. 2009; Kashiwagi et al. 2012 |
| 6 | 1754 | -7348.39 (3750.04) | -7350.44 (3753.28) | Random assignment of individuals into two species units | To assess relative support for other models. | |
| 7 | 1754 | -7360.08 (3773.42) | -7359.33 (3771.06) | Single species of manta ray (lump *M. alfredi* and *M. birostris*) | Similar levels of sequence divergence as species that have previously been lumped based on said low sequence divergence | White et al. 2017 |
| 8 | 1760 | -11864.6 (12782.46) | -11864.51 (12781.42) | Lump *M. japanica* with *M. birostris. M. alfredi* distinct | To assess evidence for interaction from higher up the tree. | |

**Supplementary Table 5**: Details of species delimitation models tested with Bayes Factor Delimitation (BFD*) within the second clade identified; *Mobula mobular* and *Mobula japanica*, including *Manta alfredi* individuals 0135, 0131, 0685 and 0146.

| Rank | SNPs retained | MLE ($2log_eBF$) - gamma prior | MLE ($2log_eBF$) - default 1/X prior | Description | Rationale | Reference |
|---|---|---|---|---|---|---|
| 1 | 1752 | -5390.81 (-119.58) | -5391.65 (-118.44) | *M. japanica* and *M. mobular* are distinct species units, where the latter is restricted to the Mediterranean Sea. | Taxonomy recognised prior to revision by White et al. 2017 | Notarbartolo di Sciara, 1987; Bustamante et al. 2016; Adnet et al, 2012 |
| 2 | 1755 | -5390.93 (-119.34) | -5393.24 (-115.26) | Split individuals into Atlantic (including Mediterranean) and Indo-Pacific species units | Distinguishability of these two groups through PCA | This study |
| 3 | 1755 | -5424.82 (-51.56) | -5426.71 (-48.32) | Random assignment of individuals into two species units | To assess relative support for other models. | |
| 4 | 1757 | -5450.6 (Null) | -5450.87 (Null) | Null model and current arrangement - *M. japanica* is a junior synonym of *M. mobular,* recognising a single species | Current taxonomic arrangement | White et al. 2017; Poortvliet et al. 2015 |
| 5 | 1759 | -9150.56 (7399.92) | -9153.48 (7405.22) | Lump *M. alfredi* with *M. mobular (*as formerly recognised; not including specimens formerly attributed to *M. japanica*) | To assess evidence for interaction from higher up the tree. | |

**Supplementary Table 6**: Details of species delimitation models tested with Bayes Factor Delimitation (BFD*) within the third clade identified; *Mobula thurstoni, Mobula kuhlii* and *Mobula eregoodootenkee*, including *Mobula hypostoma* individuals 0874, 0933, 0924 and 0992.

| Rank | SNPs retained | MLE ($2log_eBF$) - gamma prior | MLE ($2log_eBF$) - default 1/X prior | Description | Rationale | Reference |
|---|---|---|---|---|---|---|
| 1 | 1710 | -3160.68 (-1263.8) | -3159.35 (-1270) | Split *M. kuhlii* into two species units in East and West Indian Ocean, and recognise *M. thurstoni* and *M. eregoodootenkee* | Monophyly of *M. kuhlii* individuals based on geographical location sampled, and distinguishability through PCA. Monophyly and distinguishability of *M. thurstoni* and *M. eregoodootenkee* | This study |
| 2 | 1717 | -3289.06 (-1007.04) | -3291.85 (-1005) | *M. eregoodootenkee, M. kuhlii* and *M. thurstoni* are distinct species | Taxonomy recognised prior to revision by White et al. 2017 | Notarbartolo di Sciara, 1987 |
| 3 | 1731 | -3792.58 (Null) | -3794.35 (Null) | Null model and current arrangement – *M. eregoodootenkee* is a junior synonym of *M. kuhlii. M. thurstoni* is distinct | Current taxonomic arrangement | White et al. 2017 |
| 4 | 1756 | -5679.77 (3774.38) | -5683.06 (3777.42) | Random assignment of individuals to 3 species units | To assess relative support for other models. | |
| 5 | 1759 | -5818.44 (4051.72) | -5816.29 (4043.88) | Single species within this clade (lump *M. thurstoni*, *M. kuhlii* and *M. eregoodootenkee*) | For completeness | |

| | | | | | |
|---|---|---|---|---|---|
| 6 | 1733 | -8422.04 (9258.92) | -8424.08 (9259.46) | Lump *M. hypostoma* with *M. thurstoni. M. eregoodootenkee* and *M. kuhlii* distinct. | To assess evidence for interaction from higher up the tree. |

**Supplementary Table 7**: Details of species delimitation models tested with Bayes Factor Delimitation (BFD*) within the fourth clade identified; *Mobula hypostoma* and *Mobula munkiana*, including *Mobula tarapacana* individuals 0086, 0408, 0411 and 0847.

| Rank | SNPs retained | MLE ($2\log_e$BF) - gamma prior | MLE ($2\log_e$BF) - default 1/X prior | Description | Rationale | Reference |
|---|---|---|---|---|---|---|
| 1 | 1707 | -1881.75 (Null) | -1883.71 (Null) | Null model and current arranagement - *M. munkiana* and *M. hypostoma* are distinct species | Current taxonomic arrangement | Notarbartolo di Sciara, 1987; this study |
| 2 | 1731 | -2998.01 (2232.52) | -2997.63 (2227.84) | Random assignment of individuals into two species units | To assess relative support for the other models. | |
| 3 | 1731 | -3005.93 (2248.36) | -3004.82 (2242.22) | Single species within this clade - *M. munkiana* is a junior synonym of *M. hypostoma* | Suggested as a possible line of investigation in White et al. 2017 | Suggested in White et al. 2017 |
| 4 | 1748 | -6236.55 (8709.6) | -6238.57 (8709.72) | Lump *M. tarapacana* with *M. munkiana. M. hypostoma* distinct | To assess evidence for interaction from higher up the tree. | |

**Supplementary Table 8**: Individuals included in independent runs of SNAPP for species tree inference.

| Species | Sample | | | |
|---|---|---|---|---|
| | Subset 1 | Subset 2 | Subset 3 | Subset 4 |
| *M. alfredi* | 0135 | 0130 | 0132 | 0136 |
| | 0145 | 0140 | 0149 | 0146 |
| | 0688 | 0686 | 0687 | 0685 |
| *M. birostris* | 0736 | 1114 | 0732 | 0731 |
| | 0987 | 0988 | 0985 | 1110 |
| | 1168 | 1168 | 1122 | 1168 |
| *Mobula* sp. 1 | 0980 | 1241 | 0981 | 0982 |
| | 0984 | 1242 | 1240 | 0983 |
| | 1327 | 1327 | 1327 | 1239 |
| *M. eregoodootenkee* | 0684 | 0684 | 0684 | 0684 |
| | 0696 | 0696 | 0697 | 0697 |
| | 0810 | 0810 | 0813 | 0813 |
| *M. hypostoma* | 0886 | 0873 | 0924 | 0933 |
| | 0888 | 0990 | 0938 | 0991 |
| | 0993 | 0992 | 0990 | 0993 |
| *M. kuhlii* | 0524 | 0524 | 0524 | 0524 |
| | 0677 | 0582 | 0677 | 0605 |
| | 0774 | 0778 | 0777 | 0776 |
| *M. mobular* (and *M. mobular cf japanica*) | 0112 | 0104 | 0711 | 0862 |
| | 0003 | 0771 | 0101 | 0024 |
| | 0793 | 0343 | 0219 | 0773 |
| *M. munkiana* | 0821 | 0824 | 0827 | 0833 |
| | 0844 | 0844 | 0844 | 0844 |
| | 1006 | 1019 | 1006 | 1009 |
| *M. tarapacana* | 0086 | 0086 | 0086 | 0086 |
| | 0847 | 0408 | 0411 | 0411 |
| | 0408 | 0847 | 0847 | 0847 |
| *M. thurstoni* | 0159 | 0159 | 0159 | 0159 |
| | 0712 | 0712 | 0712 | 0712 |
| | 0782 | 0787 | 0786 | 0785 |
| No. SNPs retained | 1242 | 1240 | 1253 | 1250 |

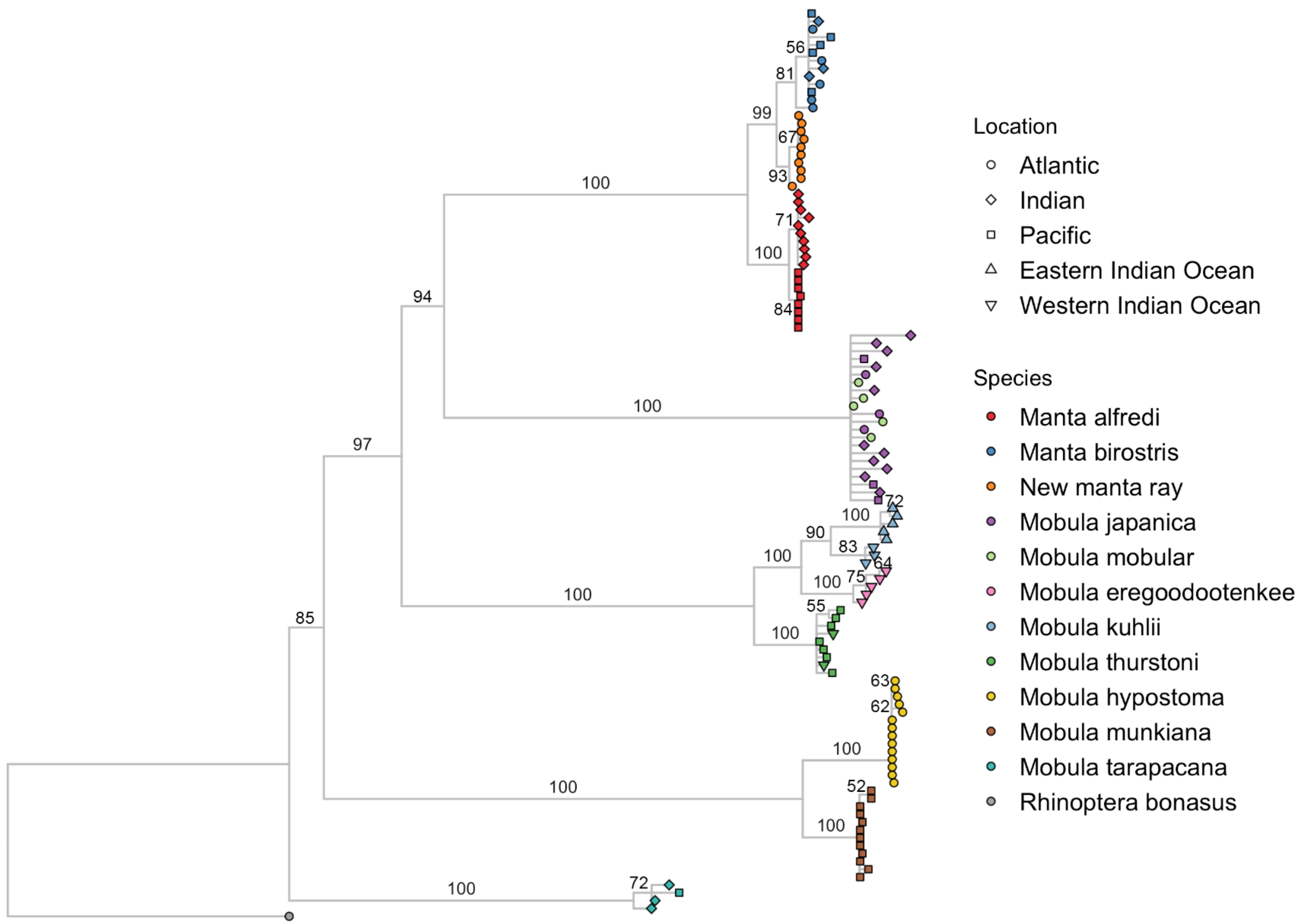

**Supplementary Figure 1**: Maximum Likelihood tree based on 1762 SNPs (dataset p90). Coloured points indicate putative species, and shape indicates geographic origin of samples as specified in the key. Bootstrap values are shown on the branches and nodes with less than 50% support are collapsed. Species names are those assigned at collection, some now considered invalid (White et al. 2017).

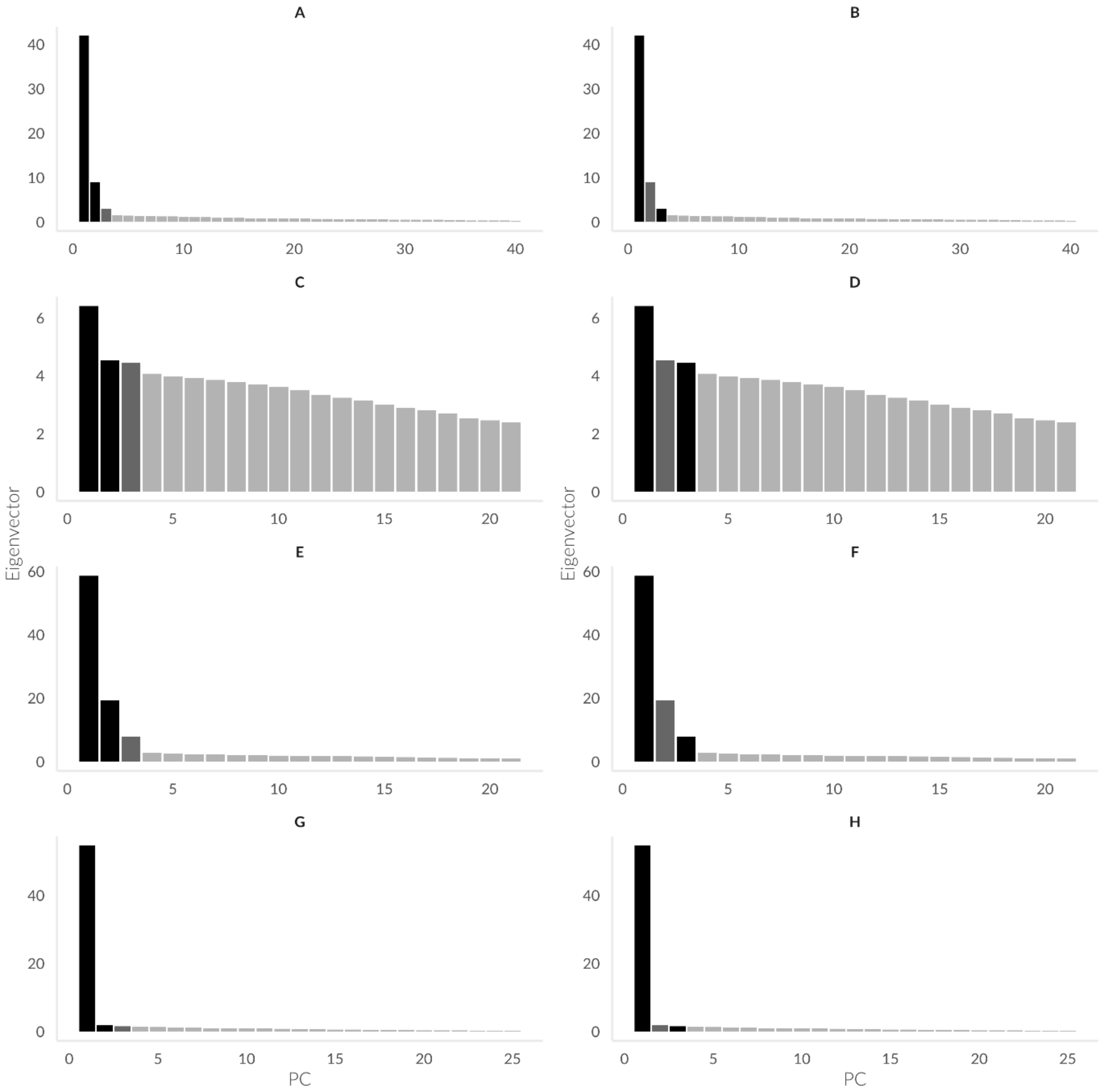

**Supplementary Figure 2**: Eigenvector plots for each Principle Components Analysis. Panel letters (A-H) correspond to the panel letters in Fig. 3. Plotted axes are in black and retained axes in dark grey.

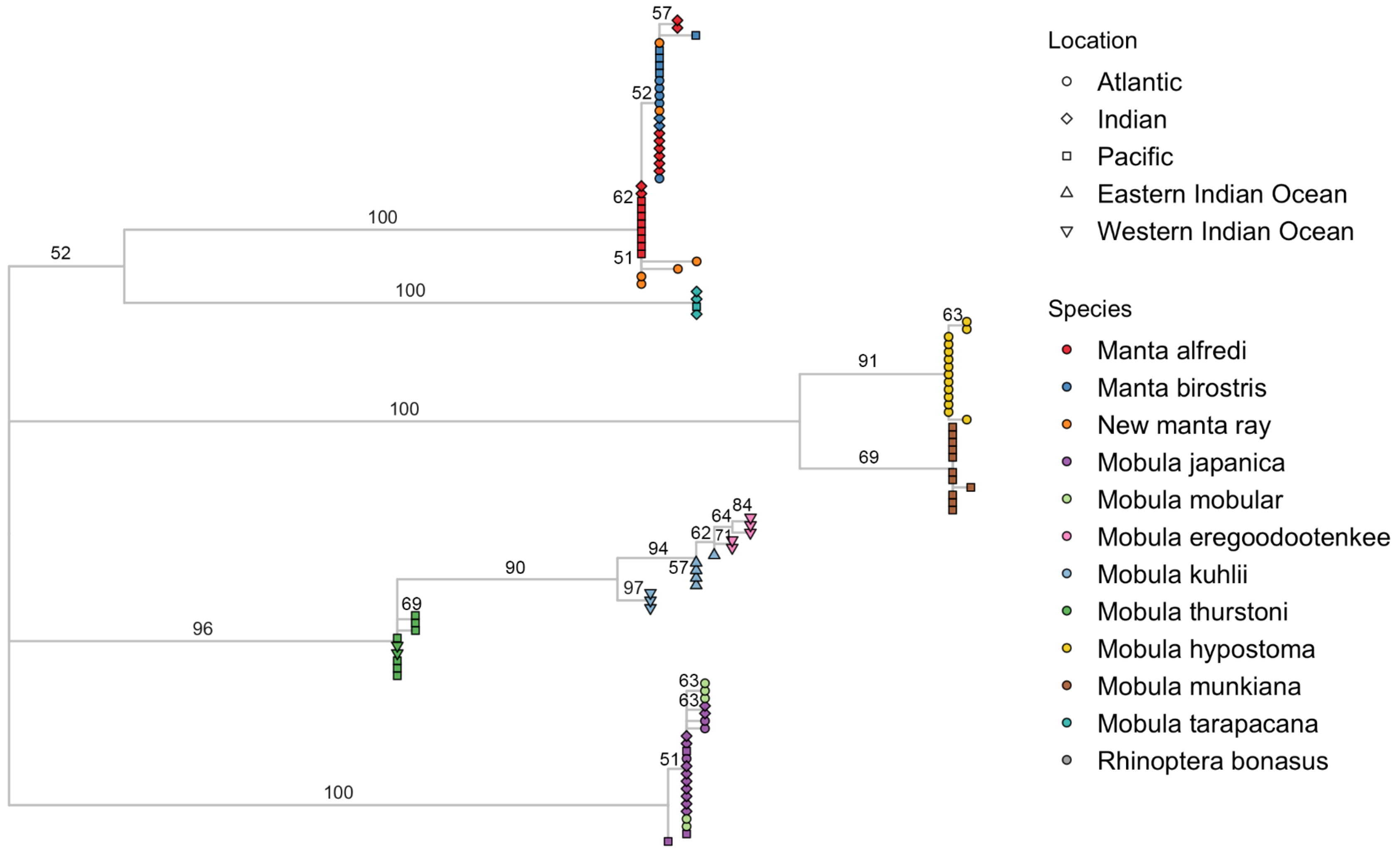

**Supplementary Figure 3**: Maximum Likelihood tree based on Cytochrome Oxidase Subunit I (COI) sequence data. Coloured points indicate putative species, and shape indicates geographic origin of samples as specified in the key. Bootstrap values are shown on the branches and nodes with less than 50% support are collapsed. Species names are those assigned to samples at collection, some now considered invalid (White et al. 2017).

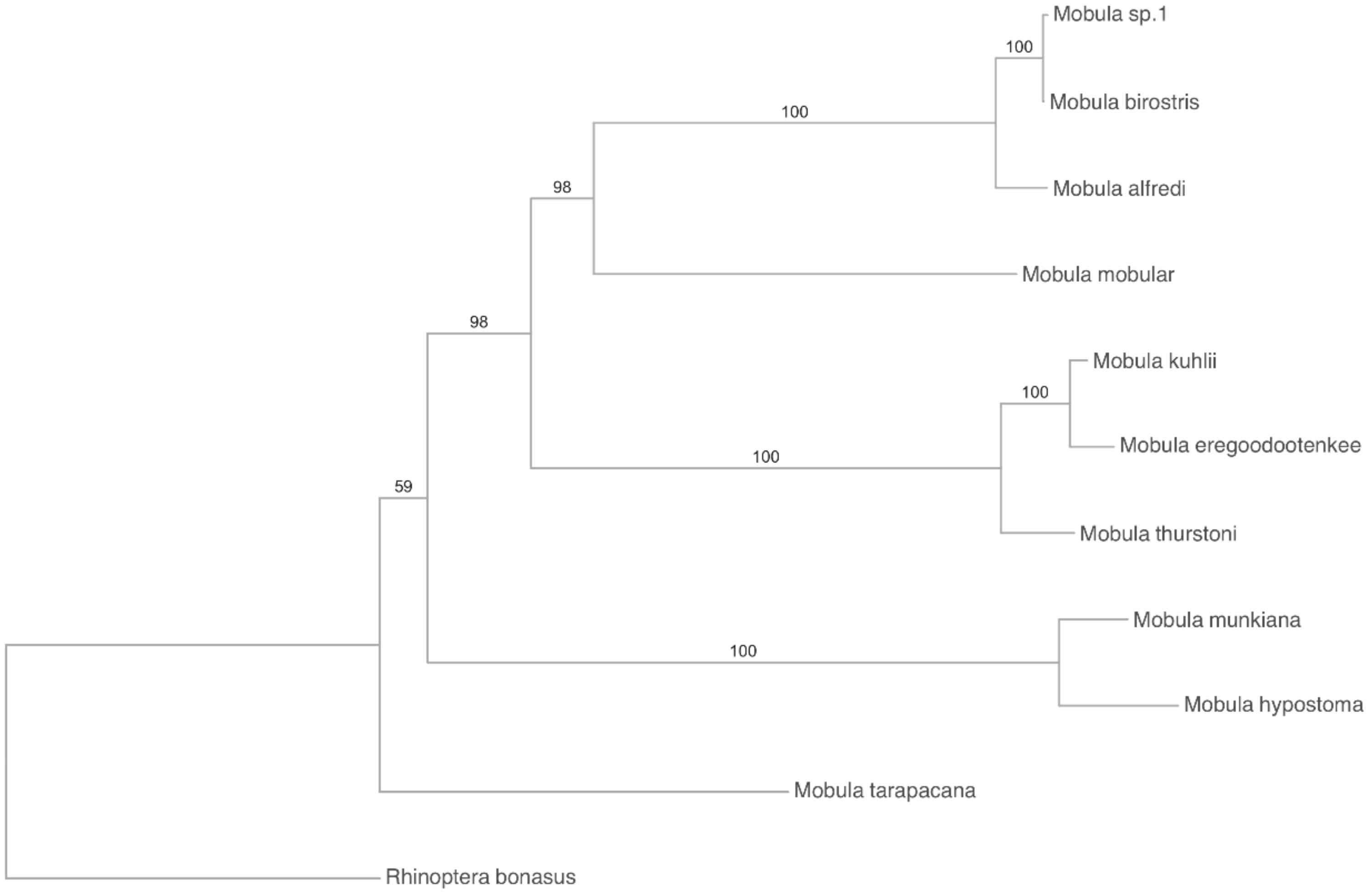

**Supplementary Figure 4**: Maximum Likelihood tree of inferred mobulid species units based on 1755 SNPs (dataset p90). Bootstrap values are shown on the branches.

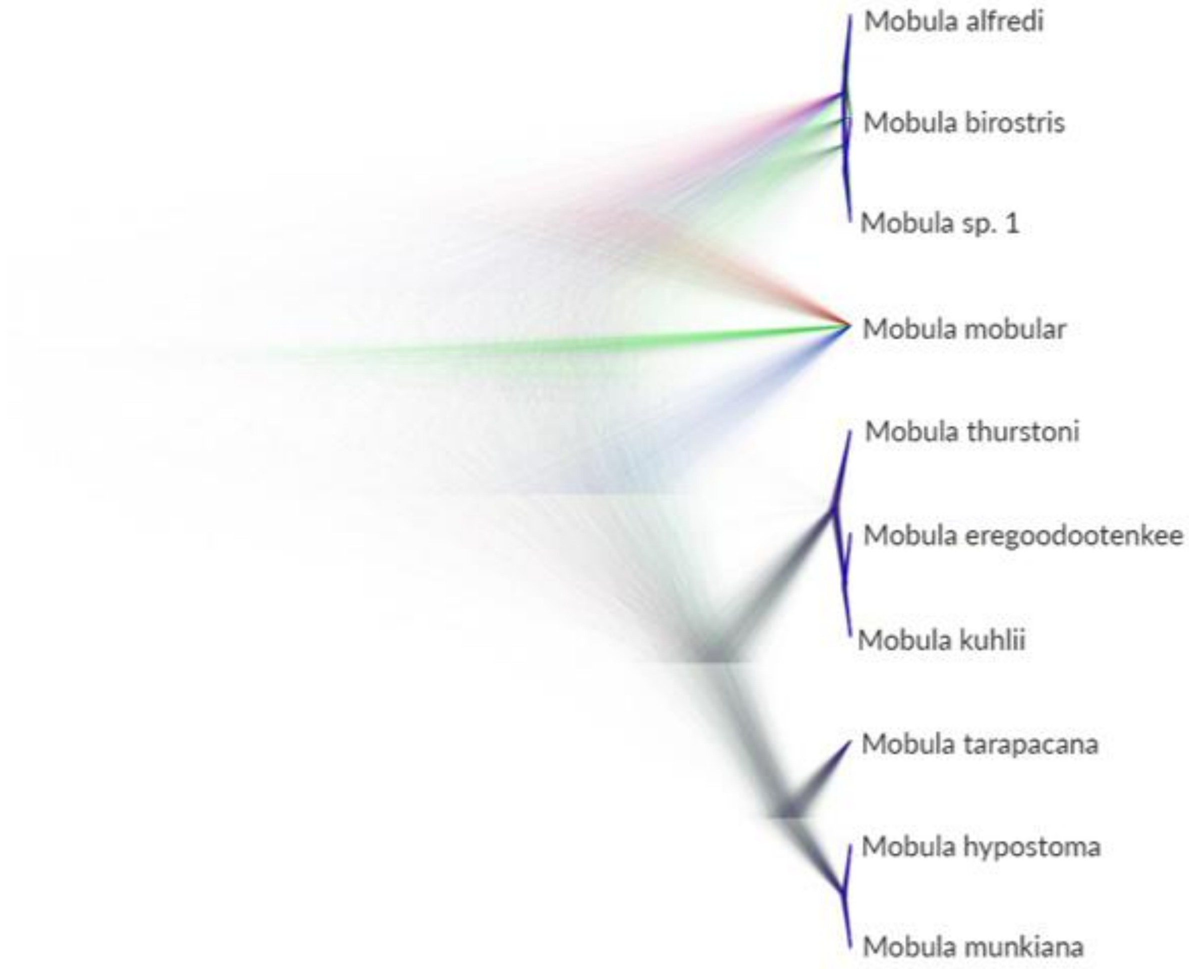

**Supplementary Figure 5**: SNP phylogeny of 30 individuals assigned to ten species based on 1240 SNPs (dataset p90, individual subset 2; Appendix S9). Tree cloud of sampled trees produced using DENSITREE (representing samples taken every 1000 MCMC steps from 5,000,0000 iterations) from SNAPP analysis to visualise the range of alternative topologies.

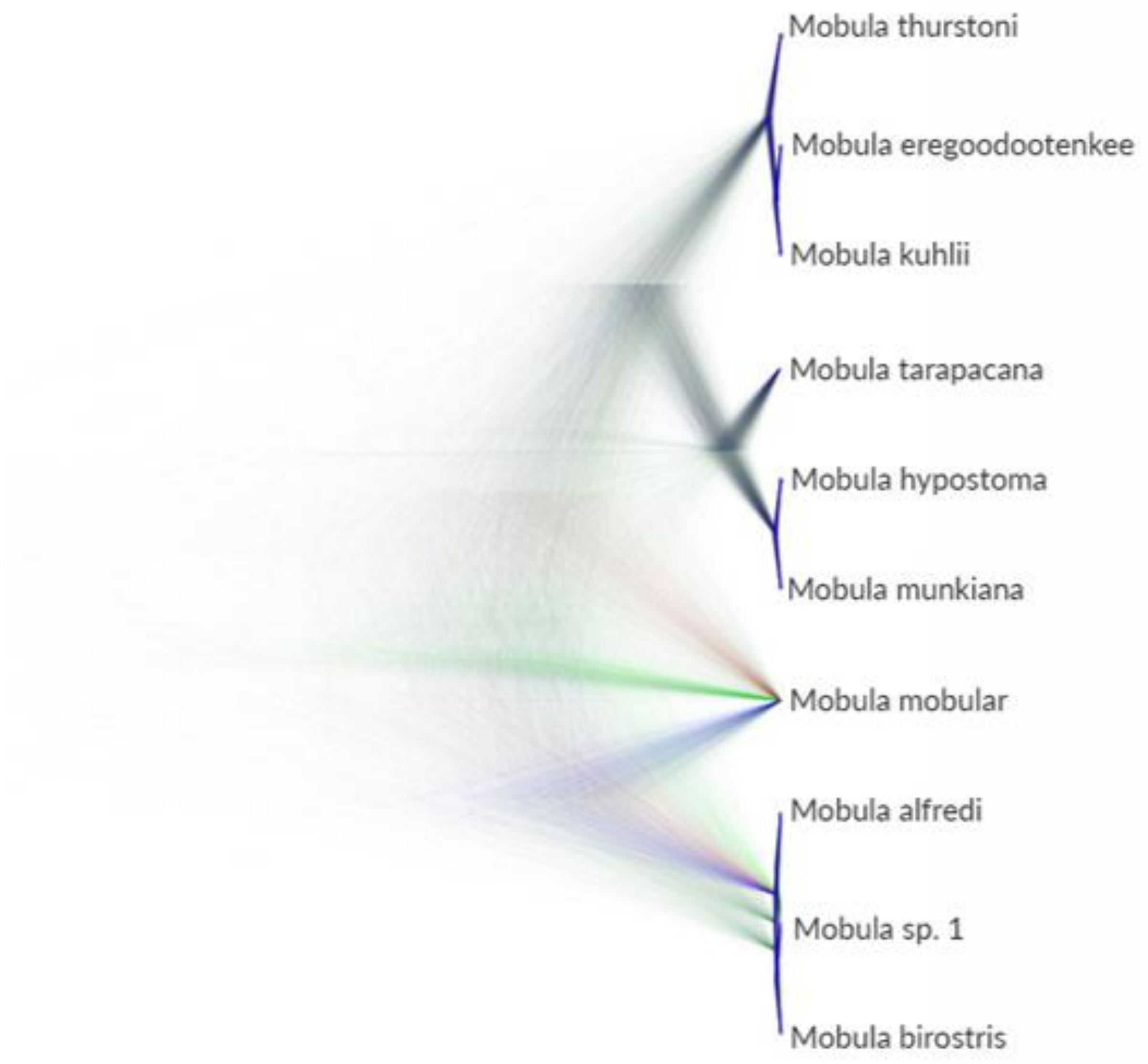

**Supplementary Figure 6**: SNP phylogeny of 30 individuals assigned to ten species based on 1253 SNPs (dataset p90, individual subset 3; Appendix S9). Tree cloud of sampled trees produced using DENSITREE (representing samples taken every 1000 MCMC steps from 5,000,0000 iterations) from SNAPP analysis to visualise the range of alternative topologies.

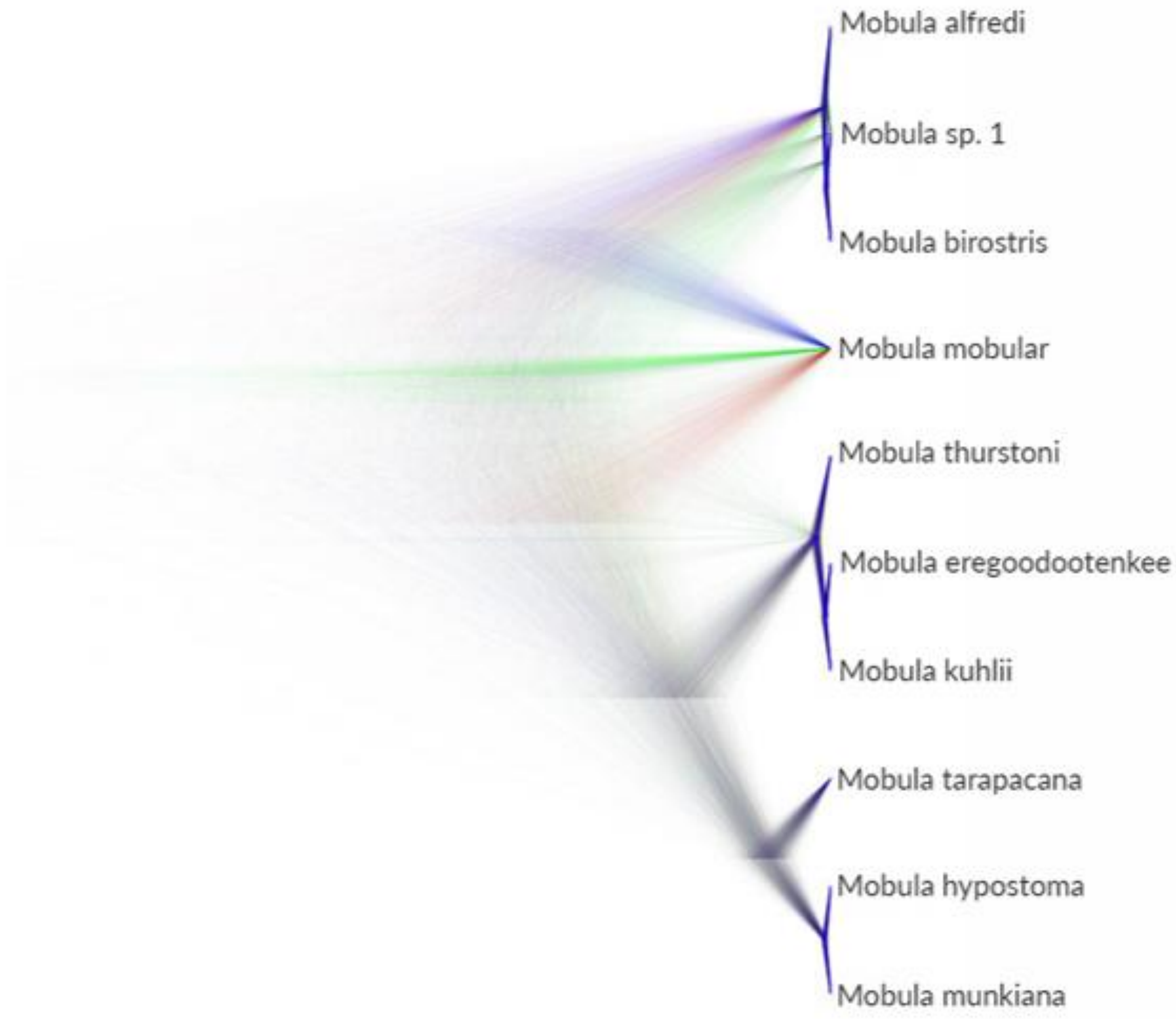

**Supplementary Figure 7**: SNP phylogeny of 30 individuals assigned to ten species based on 1250 SNPs (dataset p90, individual subset 4; Appendix S9). Tree cloud of sampled trees produced using DENSITREE (representing samples taken every 1000 MCMC steps from 5,000,0000 iterations) from SNAPP analysis to visualise the range of alternative topologies.

**Supplementary Table 9**: Topologies contained within the 95% Highest Posterior Density for each SNAPP subsample. Subsample 3 has 25 trees contained within the 95% HPD, and the first 20 are shown.

| Subsample | Tree | Percentage | Tree topology |
|---|---|---|---|
| 1 | 1 | 28.27% | *((Mobula mobular,(Mobula alfredi,(Mobula birostris,Mobula sp. 1))),((Mobula tarapacana,(Mobula munkiana,Mobula hypostoma)),(Mobula thurstoni,(Mobula kuhlii,Mobula eregoodootenkee))))* |
| | 2 | 25.62% | *((Mobula mobular,((Mobula tarapacana,(Mobula munkiana,Mobula hypostoma)),(Mobula thurstoni,(Mobula kuhlii,Mobula eregoodootenkee)))),(Mobula alfredi,(Mobula birostris,Mobula sp. 1)))* |
| | 3 | 18.52% | *(Mobula mobular,(((Mobula tarapacana,(Mobula munkiana,Mobula hypostoma)),(Mobula thurstoni,(Mobula kuhlii,Mobula eregoodootenkee))),(Mobula alfredi,(Mobula birostris,Mobula sp. 1))))* |
| | 4 | 5.0% | *((Mobula mobular,((Mobula alfredi,Mobula sp. 1),Mobula birostris)),((Mobula tarapacana,(Mobula munkiana,Mobula hypostoma)),(Mobula thurstoni,(Mobula kuhlii,Mobula eregoodootenkee))))* |
| | 5 | 4.92% | *((Mobula mobular,((Mobula tarapacana,(Mobula munkiana,Mobula hypostoma)),(Mobula thurstoni,(Mobula kuhlii,Mobula eregoodootenkee)))),((Mobula alfredi,Mobula sp. 1),Mobula birostris))* |
| | 6 | 3.92% | *((Mobula mobular,((Mobula tarapacana,(Mobula munkiana,Mobula hypostoma)),(Mobula thurstoni,(Mobula kuhlii,Mobula eregoodootenkee)))),((Mobula alfredi,Mobula birostris),Mobula sp. 1))* |
| | 7 | 3.50% | *((Mobula mobular,((Mobula alfredi,Mobula birostris),Mobula sp. 1)),((Mobula tarapacana,(Mobula munkiana,Mobula hypostoma)),(Mobula thurstoni,(Mobula kuhlii,Mobula eregoodootenkee))))* |
| | 8 | 3.47% | *(Mobula mobular,(((Mobula tarapacana,(Mobula munkiana,Mobula hypostoma)),(Mobula thurstoni,(Mobula kuhlii,Mobula eregoodootenkee))),((Mobula alfredi,Mobula sp. 1),Mobula birostris)))* |
| | 9 | 2.87% | *(Mobula mobular,(((Mobula tarapacana,(Mobula munkiana,Mobula hypostoma)),(Mobula thurstoni,(Mobula kuhlii,Mobula eregoodootenkee))),((Mobula alfredi,Mobula birostris),Mobula sp. 1)))* |

| | | | |
|---|---|---|---|
| 2 | 1 | 19.55% | *(((Mobula tarapacana,(Mobula munkiana,Mobula hypostoma)),(Mobula thurstoni,(Mobula kuhlii,Mobula eregoodootenkee))),(Mobula mobular,(Mobula alfredi,(Mobula birostris,Mobula sp. 1))))* |
| | 2 | 19.35% | *((((Mobula tarapacana,(Mobula munkiana,Mobula hypostoma)),(Mobula thurstoni,(Mobula kuhlii,Mobula eregoodootenkee))),Mobula mobular),(Mobula alfredi,(Mobula birostris,Mobula sp. 1)))* |
| | 3 | 13.17% | *((((Mobula tarapacana,(Mobula munkiana,Mobula hypostoma)),(Mobula thurstoni,(Mobula kuhlii,Mobula eregoodootenkee))),(Mobula alfredi,(Mobula birostris,Mobula sp. 1))),Mobula mobular)* |
| | 4 | 9.27% | *((((Mobula tarapacana,(Mobula munkiana,Mobula hypostoma)),(Mobula thurstoni,(Mobula kuhlii,Mobula eregoodootenkee))),Mobula mobular),((Mobula alfredi,Mobula sp. 1),Mobula birostris))* |
| | 5 | 8.75% | *(((Mobula tarapacana,(Mobula munkiana,Mobula hypostoma)),(Mobula thurstoni,(Mobula kuhlii,Mobula eregoodootenkee))),(Mobula mobular,((Mobula alfredi,Mobula sp. 1),Mobula birostris)))* |
| | 6 | 8.20% | *((((Mobula tarapacana,(Mobula munkiana,Mobula hypostoma)),(Mobula thurstoni,(Mobula kuhlii,Mobula eregoodootenkee))),Mobula mobular),((Mobula alfredi,Mobula birostris),Mobula sp. 1))* |
| | 7 | 8.05% | *(((Mobula tarapacana,(Mobula munkiana,Mobula hypostoma)),(Mobula thurstoni,(Mobula kuhlii,Mobula eregoodootenkee))),(Mobula mobular,((Mobula alfredi,Mobula birostris),Mobula sp. 1)))* |
| | 8 | 5.97% | *((((Mobula tarapacana,(Mobula munkiana,Mobula hypostoma)),(Mobula thurstoni,(Mobula kuhlii,Mobula eregoodootenkee))),((Mobula alfredi,Mobula sp. 1),Mobula birostris)),Mobula mobular)* |
| | 9 | 5.05% | *((((Mobula tarapacana,(Mobula munkiana,Mobula hypostoma)),(Mobula thurstoni,(Mobula kuhlii,Mobula eregoodootenkee))),((Mobula alfredi,Mobula birostris),Mobula sp. 1)),Mobula mobular)* |
| 3 | 1 | 18.77% | *(((Mobula tarapacana,(Mobula munkiana,Mobula hypostoma)),(Mobula thurstoni,(Mobula kuhlii,Mobula eregoodootenkee))),(Mobula mobular,(Mobula alfredi,(Mobula birostris,Mobula sp. 1))))* |
| | 2 | 14.40% | *((((Mobula tarapacana,(Mobula munkiana,Mobula hypostoma)),(Mobula thurstoni,(Mobula kuhlii,Mobula eregoodootenkee))),Mobula mobular),(Mobula alfredi,(Mobula birostris,Mobula sp. 1)))* |
| | 3 | 9.70% | *((((Mobula tarapacana,(Mobula munkiana,Mobula hypostoma)),(Mobula thurstoni,(Mobula kuhlii,Mobula eregoodootenkee))),(Mobula alfredi,(Mobula birostris,Mobula sp. 1))),Mobula mobular)* |

| | | |
|---|---|---|
| 4 | 9.25% | *(((Mobula tarapacana,(Mobula munkiana,Mobula hypostoma)),(Mobula thurstoni,(Mobula kuhlii,Mobula eregoodootenkee))),(Mobula mobular,((Mobula alfredi,Mobula sp. 1),Mobula birostris)))* |
| 5 | 7.72% | *(((Mobula tarapacana,(Mobula munkiana,Mobula hypostoma)),(Mobula thurstoni,(Mobula kuhlii,Mobula eregoodootenkee))),(Mobula mobular,((Mobula alfredi,Mobula birostris),Mobula sp. 1)))* |
| 6 | 7.0% | *((((Mobula tarapacana,(Mobula munkiana,Mobula hypostoma)),(Mobula thurstoni,(Mobula kuhlii,Mobula eregoodootenkee))),Mobula mobular),((Mobula alfredi,Mobula sp. 1),Mobula birostris))* |
| 7 | 6.37% | *((((Mobula tarapacana,(Mobula munkiana,Mobula hypostoma)),(Mobula thurstoni,(Mobula kuhlii,Mobula eregoodootenkee))),Mobula mobular),((Mobula alfredi,Mobula birostris),Mobula sp. 1))* |
| 8 | 6.20% | *((((Mobula tarapacana,(Mobula munkiana,Mobula hypostoma)),(Mobula thurstoni,(Mobula kuhlii,Mobula eregoodootenkee))),((Mobula alfredi,Mobula sp. 1),Mobula birostris)),Mobula mobular)* |
| 9 | 4.90% | *((((Mobula tarapacana,(Mobula munkiana,Mobula hypostoma)),(Mobula thurstoni,(Mobula kuhlii,Mobula eregoodootenkee))),((Mobula alfredi,Mobula birostris),Mobula sp. 1)),Mobula mobular)* |
| 10 | 1.12% | *(((Mobula tarapacana,(Mobula munkiana,Mobula hypostoma)),(Mobula mobular,(Mobula thurstoni,(Mobula kuhlii,Mobula eregoodootenkee)))),(Mobula alfredi,(Mobula birostris,Mobula sp. 1)))* |
| 11 | 0.92% | *(((Mobula tarapacana,(Mobula munkiana,Mobula hypostoma)),Mobula mobular),((Mobula alfredi,(Mobula birostris,Mobula sp. 1)),(Mobula thurstoni,(Mobula kuhlii,Mobula eregoodootenkee))))* |
| 12 | 0.90% | *((((Mobula tarapacana,(Mobula munkiana,Mobula hypostoma)),Mobula mobular),(Mobula thurstoni,(Mobula kuhlii,Mobula eregoodootenkee))),(Mobula alfredi,(Mobula birostris,Mobula sp. 1)))* |
| 13 | 0.90% | *(((Mobula tarapacana,(Mobula munkiana,Mobula hypostoma)),(Mobula alfredi,(Mobula birostris,Mobula sp. 1))),(Mobula mobular,(Mobula thurstoni,(Mobula kuhlii,Mobula eregoodootenkee))))* |
| 14 | 0.72% | *(((Mobula tarapacana,(Mobula munkiana,Mobula hypostoma)),(Mobula mobular,(Mobula thurstoni,(Mobula kuhlii,Mobula eregoodootenkee)))),((Mobula alfredi,Mobula sp. 1),Mobula birostris))* |
| 15 | 0.72% | *((((Mobula tarapacana,(Mobula munkiana,Mobula hypostoma)),(Mobula alfredi,(Mobula birostris,Mobula sp. 1))),(Mobula thurstoni,(Mobula kuhlii,Mobula eregoodootenkee))),Mobula mobular)* |

| | | | |
|---|---|---|---|
| | 16 | 0.70% | *(((Mobula tarapacana,(Mobula munkiana,Mobula hypostoma)),((Mobula alfredi,Mobula sp. 1),Mobula birostris)),(Mobula mobular,(Mobula thurstoni,(Mobula kuhlii,Mobula eregoodootenkee))))* |
| | 17 | 0.62% | *(((Mobula tarapacana,(Mobula munkiana,Mobula hypostoma)),((Mobula alfredi,(Mobula birostris,Mobula sp. 1)),(Mobula thurstoni,(Mobula kuhlii,Mobula eregoodootenkee)))),Mobula mobular)* |
| | 18 | 0.57% | *((((Mobula tarapacana,(Mobula munkiana,Mobula hypostoma)),Mobula mobular),(Mobula thurstoni,(Mobula kuhlii,Mobula eregoodootenkee))),((Mobula alfredi,Mobula sp. 1),Mobula birostris))* |
| | 19 | 0.55% | *((((Mobula tarapacana,(Mobula munkiana,Mobula hypostoma)),Mobula mobular),(Mobula alfredi,(Mobula birostris,Mobula sp. 1))),(Mobula thurstoni,(Mobula kuhlii,Mobula eregoodootenkee)))* |
| | 20 | 0.52% | *((Mobula tarapacana,(Mobula munkiana,Mobula hypostoma)),((Mobula mobular,(Mobula alfredi,(Mobula birostris,Mobula sp. 1))),(Mobula thurstoni,(Mobula kuhlii,Mobula eregoodootenkee))))* |
| 4 | 1 | 23.95% | *((Mobula mobular,(Mobula alfredi,(Mobula birostris,Mobula sp. 1))),((Mobula tarapacana,(Mobula munkiana,Mobula hypostoma)),(Mobula thurstoni,(Mobula kuhlii,Mobula eregoodootenkee))))* |
| | 2 | 21.84% | *((Mobula mobular,((Mobula tarapacana,(Mobula munkiana,Mobula hypostoma)),(Mobula thurstoni,(Mobula kuhlii,Mobula eregoodootenkee)))),(Mobula alfredi,(Mobula birostris,Mobula sp. 1)))* |
| | 3 | 15.97% | *(Mobula mobular,(((Mobula tarapacana,(Mobula munkiana,Mobula hypostoma)),(Mobula thurstoni,(Mobula kuhlii,Mobula eregoodootenkee))),(Mobula alfredi,(Mobula birostris,Mobula sp. 1))))* |
| | 4 | 5.40% | *((Mobula mobular,((Mobula alfredi,Mobula sp. 1),Mobula birostris)),((Mobula tarapacana,(Mobula munkiana,Mobula hypostoma)),(Mobula thurstoni,(Mobula kuhlii,Mobula eregoodootenkee))))* |
| | 5 | 5.30% | *((Mobula mobular,((Mobula tarapacana,(Mobula munkiana,Mobula hypostoma)),(Mobula thurstoni,(Mobula kuhlii,Mobula eregoodootenkee)))),((Mobula alfredi,Mobula sp. 1),Mobula birostris))* |
| | 6 | 4.82% | *((Mobula mobular,((Mobula alfredi,Mobula birostris),Mobula sp. 1)),((Mobula tarapacana,(Mobula munkiana,Mobula hypostoma)),(Mobula thurstoni,(Mobula kuhlii,Mobula eregoodootenkee))))* |
| | 7 | 4.70% | *((Mobula mobular,((Mobula tarapacana,(Mobula munkiana,Mobula hypostoma)),(Mobula thurstoni,(Mobula kuhlii,Mobula eregoodootenkee)))),((Mobula alfredi,Mobula birostris),Mobula sp. 1))* |

| | | |
|---|---|---|
| 8 | 3.57% | *(Mobula mobular,(((Mobula tarapacana,(Mobula munkiana,Mobula hypostoma)),(Mobula thurstoni,(Mobula kuhlii,Mobula eregoodootenkee))),((Mobula alfredi,Mobula sp. 1),Mobula birostris)))* |
| 9 | 2.85% | *(Mobula mobular,(((Mobula tarapacana,(Mobula munkiana,Mobula hypostoma)),(Mobula thurstoni,(Mobula kuhlii,Mobula eregoodootenkee))),((Mobula alfredi,Mobula birostris),Mobula sp. 1)))* |
| 10 | 1.37% | *(((Mobula mobular,(Mobula thurstoni,(Mobula kuhlii,Mobula eregoodootenkee))),(Mobula tarapacana,(Mobula munkiana,Mobula hypostoma))),(Mobula alfredi,(Mobula birostris,Mobula sp. 1)))* |
| 11 | 1.12% | *(((Mobula mobular,(Mobula tarapacana,(Mobula munkiana,Mobula hypostoma))),(Mobula thurstoni,(Mobula kuhlii,Mobula eregoodootenkee))),(Mobula alfredi,(Mobula birostris,Mobula sp. 1)))* |
| 12 | 0.75% | *((Mobula mobular,(Mobula thurstoni,(Mobula kuhlii,Mobula eregoodootenkee))),((Mobula tarapacana,(Mobula munkiana,Mobula hypostoma)),(Mobula alfredi,(Mobula birostris,Mobula sp. 1))))* |
| 13 | 0.72% | *((Mobula mobular,(Mobula tarapacana,(Mobula munkiana,Mobula hypostoma))),((Mobula alfredi,(Mobula birostris,Mobula sp. 1)),(Mobula thurstoni,(Mobula kuhlii,Mobula eregoodootenkee))))* |
| 14 | 0.70% | *(Mobula mobular,((Mobula tarapacana,(Mobula munkiana,Mobula hypostoma)),((Mobula alfredi,(Mobula birostris,Mobula sp. 1)),(Mobula thurstoni,(Mobula kuhlii,Mobula eregoodootenkee)))))* |
| 15 | 0.57% | *(Mobula mobular,(((Mobula tarapacana,(Mobula munkiana,Mobula hypostoma)),(Mobula alfredi,(Mobula birostris,Mobula sp. 1))),(Mobula thurstoni,(Mobula kuhlii,Mobula eregoodootenkee))))* |
| 16 | 0.52% | *(((Mobula mobular,(Mobula alfredi,(Mobula birostris,Mobula sp. 1))),(Mobula tarapacana,(Mobula munkiana,Mobula hypostoma))),(Mobula thurstoni,(Mobula kuhlii,Mobula eregoodootenkee)))* |
| 17 | 0.50% | *(((Mobula mobular,(Mobula alfredi,(Mobula birostris,Mobula sp. 1))),(Mobula thurstoni,(Mobula kuhlii,Mobula eregoodootenkee))),(Mobula tarapacana,(Mobula munkiana,Mobula hypostoma)))* |
| 18 | 0.40% | *(((Mobula mobular,(Mobula thurstoni,(Mobula kuhlii,Mobula eregoodootenkee))),(Mobula alfredi,(Mobula birostris,Mobula sp. 1))),(Mobula tarapacana,(Mobula munkiana,Mobula hypostoma)))* |

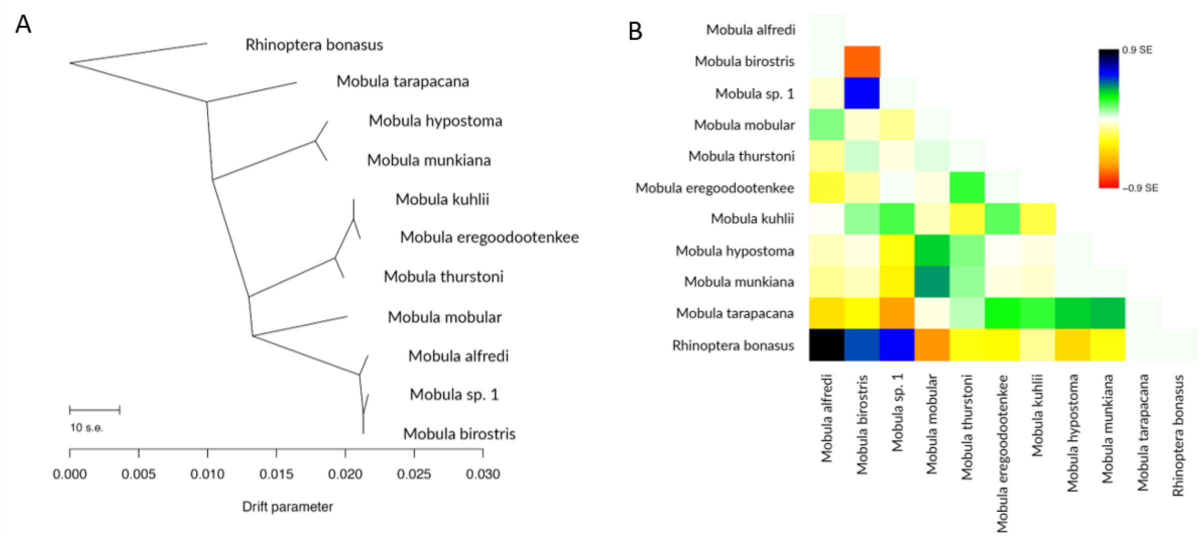

**Supplementary Figure 8**: A) Admixture graph showing relationships among inferred mobulid species units as a simple bifurcating tree, inferred using a Maximum Likelihood method in TreeMix. Horizontal branch lengths represent drift. The scale bar shows 10 times the average standard error of the values in the sample covariance matrix. This model explains 99.86% of the variance in the data. B) Residual fit of the observed versus predicted squared allele frequency difference.

**Supplementary Table 10**: f3-statistics for comparisons of different clade topologies. *M. alfredi, M. thurstoni* and *M. mobular* were randomly chosen to represent their respective clades.

| 3-taxon tree | $f_3$-statistic ± SE | Z-score |
|---|---|---|
| *M.alfredi;M.mobular,M.thurstoni* | 0.092 ± 0.005 | 17.8885 |
| *M.mobular;M.alfredi,M.thurstoni* | 0.0854984 ± 0.00475089 | 17.9963 |
| *M.thurstoni;M.alfredi,M.mobular* | 0.101701 0.00534773 | 19.0177 |

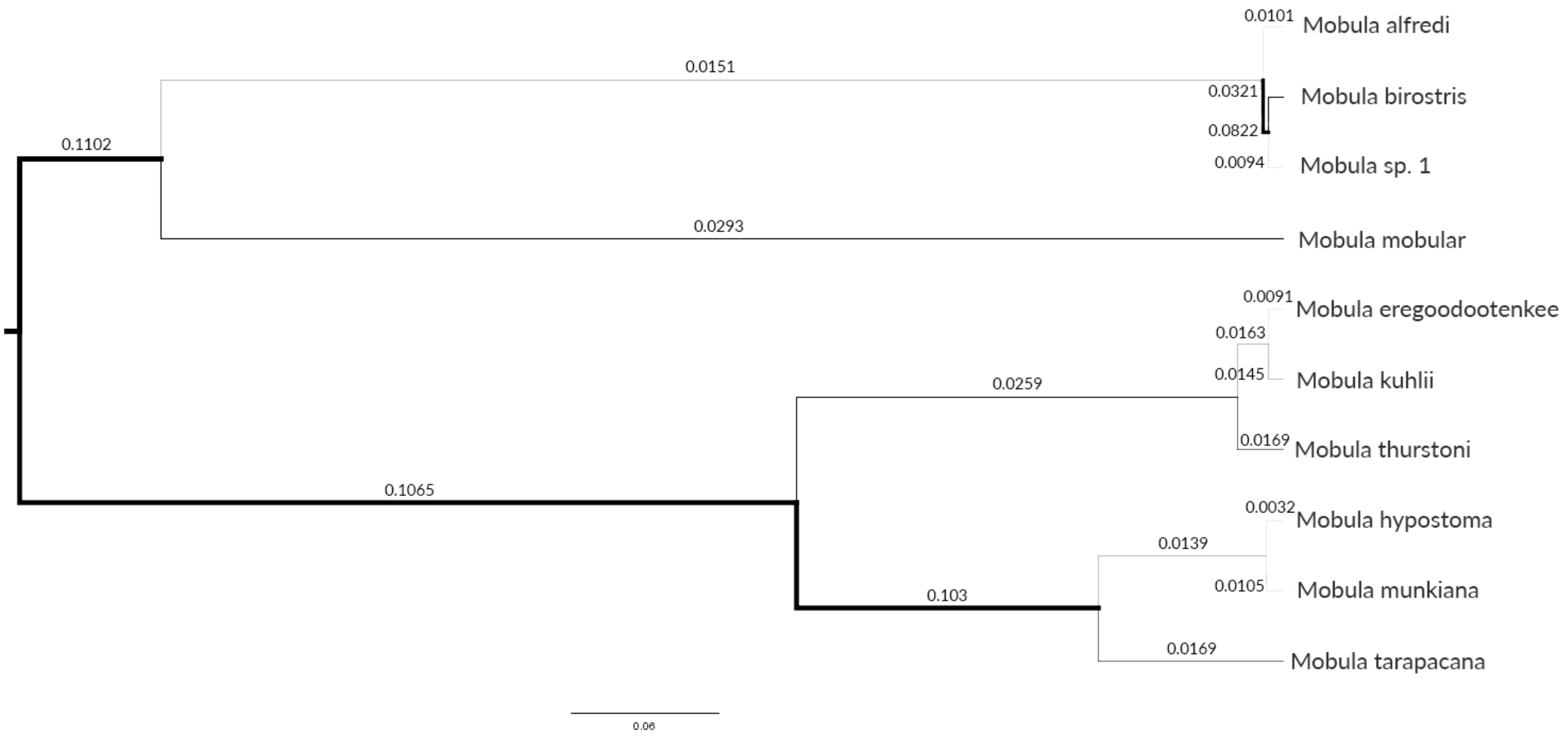

**Supplementary Figure 9**: Maximum-clade-credibility tree for inferred mobulid species units using SNAPP (individual subset 1; Appendix S9). Branch width is proportional to theta, and theta values are shown on the branches.

### Appendix 1: Taxonomic Implications

Through the analysis of genome-wide data for a globally and taxonomically comprehensive set of mobulid tissue samples, we produced the most extensive phylogeny for the Mobulidae to date, and carried out species delimitation using a multispecies coalescent based approach. As such, our findings have implications for mobulid taxonomy. It is important to recognise speciation as a continuous process, however, where lineage splitting does not necessarily correspond to speciation events. When this is explicitly modelled, the multispecies coalescent has been shown to overestimate species numbers, recovering all structure both at the level of the species and the population (Sukumaran & Knowles, 2017). In contrast to previous studies evaluating mobulid diversity, the global nature of our dataset allows for this conflict to be resolved, where in many cases, individuals from pairs of putative species are sampled within sites (Fig. 1), thereby allowing for the identification of true reproductive isolation. We summarise and discuss these taxonomic implications below since it will be of general interest to policy-makers. Additionally, it is provided as a resource for taxonomists wishing to compliment traditional approaches, such as morphological observation of specimens or ecological and behavioural data, with genomic data to evaluate taxonomy of the Mobulidae.

In brief, our genome-wide SNP data provide evidence supporting ten species within the Mobulidae. Of those species defined by White et al. (2017), our data support *Mobula alfredi, Mobula birostris, Mobula mobular, Mobula thurstoni, Mobula hypostoma, Mobula munkiana* and *Mobula tarapacana*. However, our data strongly suggests that individuals identified as *Mobula kuhlii* and *Mobula eregoodootenkee* prior to the revision published by White et al. (2017) are distinct and reproductively isolated, thereby belonging to separate species. Furthermore, we find strong evidence for a currently undescribed species of manta ray (*Mobula* sp. 1) in the Gulf of Mexico. We emphatically urge policy-makers, particularly the large conventions (such as CITES and CMS) and the relevant IUCN specialist group to evaluate these as separate units in assessments and when implementing conservation policy.

### Validity of the genus *Manta*

The two species in the genus *Manta* have recently been subsumed into *Mobula* (White et al. 2017), meaning that the names *Mobula alfredi* and *Mobula birostris* are considered valid and are in current use. Our Maximum Likelihood phylogenetic analysis indicates that the previously recognised genus *Manta* is nested within *Mobula*, and provides further justification for the associated change in nomenclature implemented by White et al. (2017). However, application of a multispecies coalescent-based approach to our data allowed visualisation of the uncertainty in species tree topology and incomplete lineage sorting. Whilst our Bayesian multispecies coalescent analyses do not specifically refute the observation that *Manta* is nested within *Mobula*, we find substantial uncertainty in the placement of *M. mobular* (Fig. 5; Appendices S14-S16). Trees within the 95% Highest Posterior Density (HPD) that place *M. mobular* with the manta rays are present in approximately equal proportions to trees placing the species with the remaining devil rays (Appendix S17), thereby producing trees where the two formerly recognised genera are reciprocally monophyletic. Our results indicate that this uncertainty can be attributed to incomplete lineage sorting rather than admixture or introgression, and standing variation in extinct ancestral populations is hypothesised to drive taxonomic uncertainty with respect to the validity of the genus *Manta* (see manuscript for discussion). Given that recently separated populations or species will pass through stages of polyphyly and paraphyly before becoming reciprocally monophyletic in the absence of additional introgression (Avise 1990; Patton & Smith, 1994), it is reasonable to hypothesise we are observing this process in the Mobulidae. Based on current information however, our data are in agreement with the conclusion of White et al. (2017) with respect to the validity of the genus *Manta*, and further support the names *Mobula alfredi* and *Mobula birostris* as valid.

*Mobula* sp. 1

We find strong evidence supporting the existence of a third, undescribed species of manta ray in the Gulf of Mexico (hereafter referred to as '*Mobula* sp. 1'). Samples were analysed from two sites within the Gulf of Mexico; offshore of the Yucatan Peninsula and Flower Garden Banks National Marine Sanctuary, and were initially identified as *Manta birostris* (now *Mobula birostris*) (Fig. 1). When these Gulf of Mexico samples were analysed alongside *M. birostris* samples collected elsewhere (Sri Lanka, the Philippines and the Pacific side of Mexico), individuals were found to fall within two distinct groups; one containing only individuals from the Gulf of Mexico sites, and the other containing additional individuals from these same Gulf of Mexico sites as well as *M. birostris* individuals sampled elsewhere (Figs. 2 & 3A). In addition, we find decisive support for models recognising these groups as distinct species through Bayes Factor Delimitation (Fig. 2). Given that samples from both groups were collected within Gulf of Mexico sites, *M. birostris* can be considered to occur in sympatry with *Mobula* sp. 1, constituting separately evolving lineages (De Queiroz, 2007). Monophyly of groups supports these as separate species under the phylogenetic species concept (Frankham et al. 2012). Furthermore, sympatry of populations suggests reproductive isolation driven either by a factor other than geographical separation, or historical separation followed by modern secondary contact (as hypothesised by Hinojosa-Alvarez et al. 2016), and these species are therefore further supported under the Biological Species concept (Frankham et al. 2012). In addition, we report on a single individual which could be considered genetically intermediate between the two groups (Figs. 2 & 3), indicating that hybridisation may occur between the two species, as between *M. alfredi* and *M. birostris* (Walter et al. 2014).

In addition to previous observations of possible morphological differences (Marshall et al. 2009), novel mitochondrial DNA haplotypes have also been reported from manta rays off the Yucatan Peninsula, and a speciation event hypothesised (Hinojosa-Alvarez et al. 2016). Our study is the first analysis of genome-wide data to suggest that there are two species of manta ray in the Gulf of Mexico; a finding that is consistent with previous studies (Hinojosa-Alvarez et al. 2016; Stewart et al. 2018a). Monophyly of groups indicate that some *M. birostris* individuals using sites in the Gulf of Mexico are more closely related to *M. birostris* in Sri Lanka and the Philippines than to individuals of *Mobula* sp. 1 using those same Gulf of Mexico sites. It is therefore likely that these species occur in a state of mosaic sympatry, as with *M. alfredi* and *M. birostris* elsewhere (Kashiwagi et al. 2011). For effective conservation it will be necessary to formally describe this new species and determine the extent of its range.

#### *Mobula kuhlii* and *Mobula eregoodootenkee*

A recent taxonomic review concluded that *Mobula eregoodootenkee* is a junior synonym of *Mobula kuhlii* based on mitogenome and nuclear data for a single sample per putative species (White et al. 2017). In direct contrast, our phylogenetic analysis of genome-wide SNPs for multiple individuals per species from a broad geographic range placed individuals of *M. kuhlii* and *M. eregoodootenkee* into discrete monophyletic clades with very high bootstrap support (Fig. 2). This pattern was also mirrored in the results of our Principal Components Analysis (PCA; Fig. 3E). Bayes Factor Delimitation models that recognised individuals identified as *M. eregoodootenkee* as a distinct species from *M. kuhlii* were consistently favoured over the null model where these individuals are considered a single species (Fig. 2). Given that both species groups included samples that were collected within the same ~120km stretch of South African coastline, the divergence reported here between *M. kuhlii* and *M. eregoodootenkee* cannot be attributed to geographic population structure (Sukumaran & Knowles, 2017). There is evidence to suggest that periods of speciation within the Mobulidae correspond to episodes of global warming and associated changes in upwelling intensity and productivity, and it is hypothesized that this led to fragmentation and subsequent divergence with respect to feeding strategies (Poortvliet et al. 2015). Differences in morphology between specimens identified as *M. kuhlii* and *M. eregoodootenkee* (Notarbartolo Di Sciara, 1987; Notarbartolo di Sciara et al. 2017), and particularly differences in the length of the cephalic fins and gill plate morphology (Paig-Tran et al.

2013) that relate directly to the filter feeding strategy of mobulid rays may lend support to this hypothesis. Notwithstanding, the present study provides robust evidence from genomic data that individuals identified as *Mobula eregoodootenkee* belong to a distinct species to individuals identified as *Mobula kuhlii* as recognised prior to White et al. (2017).

#### *Mobula mobular* and *Mobula japanica*

A recent taxonomic review concluded that *Mobula japanica* is a junior synonym of *Mobula mobular* (White et al. 2017). In agreement with this conclusion, we find no evidence from genome-wide SNPs to support *M. japanica* as a distinct species to *M. mobular*. Individuals identified as *M. mobular* as formerly recognised (with a distribution restricted to the Mediterranean Sea) do not form a reciprocally monophyletic group to the exclusion of individuals belonging to *M. japanica* (circumglobally distributed with the exception of the Mediterranean Sea), and instead these individuals form a single clade with high bootstrap support (Fig. 2). Clustering analyses indicate a degree of population structure (Fig. 3C-D), with some modest differentiation between Indo-Pacific and Atlantic (including Mediterranean) groups ($F_{ST}$ = 0.06). Results from Bayes Factor Delimitation are far less conclusive than those for other clades (Fig. 2), and support for split models being driven by geographic segregation of populations cannot be ruled out (Sukumaran & Knowles, 2017). Our data therefore supports *Mobula mobular* as a single species unit, with *M. japanica* a junior synonym.

#### *Mobula hypostoma* and *Mobula munkiana*

In their recent taxonomic review, White et al. (2017) reported a close relationship between *Mobula hypostoma* and *M. munkiana* observed through analysis of nuclear exon data of a single sample per species and commented that further research is required to ascertain whether these are truly separate species. Our genome-wide SNP data provide evidence to support *M. hypostoma* and *M. munkiana* as distinct species units (Figs. 2 & 3G-H). Whilst these species are geographically segregated in the Atlantic and Eastern Pacific Oceans respectively, our data indicates divergence of a similar magnitude to that of other distinct species groups within the Mobulidae (Figs. 2 & 3, Appendix S10).

#### *Mobula rochebrunei*

A recent taxonomic review concluded that *Mobula rochebrunei* (a pygmy devil ray species described off the coast of West Africa) is a junior synonym of *Mobula hypostoma,* based on mitogenome data for a single sample per putative species (White et al. 2017). However, mitogenome data is considered unsuitable for species delimitation or phylogenetics when used in isolation (Petit & Excoffier, 2009), and previous studies have concluded *M. rochebrunei* is a distinct species based on morphological differences (Cadenat, 1960; Notarbartolo Di Sciara, 1987). In this study, we were unable to generate molecular data representing *M. rochebrunei*, due to the only available sample being from a museum specimen stored in formalin, yielding no DNA. Notwithstanding, the revision published by White et al. (2017) is consistent with equivalent mitochondrial sequence divergence estimates for mobulid groups where further study has resolved separate species: *M. alfredi* and *M. birostris* (Marshall et al. 2009; Kashiwagi et al. 2012; this study), and *M. kuhlii* and *M. eregoodootenkee* (this study). Given the extent of illegal, unreported and unregulated (IUU) fishing pressure in the West African region (Agnew et al. 2009) and the high vulnerability to extinction which exists for mobulid species with restricted ranges (Atta-Mills et al. 2004; Doumbouya 2009) efforts to evaluate mobulid diversity in West Africa should be given a high priority (see Stewart et al. 2018b).

#### Other diversity within the Mobulidae

We identify substantial geographically-mediated population structure within *Mobula kuhlii* and *Manta alfredi* (now *Mobula alfredi*; White et al. 2017) with genome-wide SNPs. In *M. kuhlii*, individuals sampled across the Indian Ocean fall into reciprocally monophyletic groups with high bootstrap support (Fig. 2). Consistently, individuals from the East and West Indian Ocean are separated into distinct clusters through PCA (Fig. 3F), with substantial differentiation ($F_{ST}$ = 0.32). Models recognising these populations as distinct species were favoured through Bayes Factor Delimitation (Fig. 2). Indeed, there are anecdotal suggestions of morphological differences occurring in *M. kuhlii* across the Indian Ocean (Stevens et al. 2018). We identified similar patterns in *M. alfredi*. Individuals sampled in the Indian and Pacific Oceans formed distinct monophyletic groups (Fig. 2), visible as clusters through PCA (Fig. 3B) and exhibited substantial differentiation ($F_{ST}$ = 0.16). Furthermore, models recognising these populations as distinct species were favoured through Bayes Factor Delimitation (Fig. 2).

However, approaches based on the multispecies coalescent, such as Bayes Factor Delimitation, recover all structure both at the level of the species and the population (Sukumaran & Knowles, 2017). In *M. kuhlii* and *M. alfredi*, we cannot rule out a geographic driver of the patterns observed, which should therefore be tentatively attributed to population structure over speciation. Our study aimed to delimit species units for conservation and understand phylogenetic relationships. It is therefore important to recognise that the set of samples analysed here is limited to a few individuals per population, and a limited number of populations are sampled across the broad geographic ranges of these species, within which large areas are unrepresented. Detailed inferences regarding population genetic structure are therefore not possible since capturing population level diversity requires a much more comprehensive sampling regime, and results must be interpreted with caution. Further research is therefore required to determine whether the large differences within *M. kuhlii* and *M. alfredi* observed here are due to barriers to gene flow, isolation by distance across ocean clines or other factors. Nonetheless, we find sufficient intra-specific diversity to indicate potential for determining regional location of catch in these species, and indeed this is increasingly required to comply with global requirements to include capture location and species in trade (Nielsen et al. 2012). Such further study to assess population genetic structure of both species, and other species of mobulid ray, would be prudent to support effective management.

Notarbartolo Di Sciara G. 1987. A revisionary study of the genus Mobula Rafinesque, 1810 (Chondrichthyes: Mobulidae) with the description of a new species. Zoological Journal of the Linnean

Society **91**(1):1–91.

Paig-Tran EWM, Kleinteich T and Summers AP. 2013. The filter pads and filtration mechanisms of the devil rays: Variation at macro and microscopic scales. Journal of Morphology **274**(9):1026–1043.

Patton JL and Smith MF. 1994. Paraphyly, polyphyly, and the nature of species boundaries in pocket gophers (genus thomomys). Systematic Biology **43**(1):11-26.

Poortvliet M, et al. 2015. A dated molecular phylogeny of manta and devil rays (Mobulidae) based on mitogenome and nuclear sequences. Molecular Phylogenetics and Evolution **83**:72–85.

De Queiroz K. 2007. Species concepts and species delimitation. Systematic Biology **56**(6):879–886.

Stevens G, Fernando D, Dando M, Notarbartolo Di Sciara GN. 2018. Guide to the Manta and Devil Rays of the World. Princeton University Press.

Stewart JD, Nuttall M, Hickerson EL and Johnston MA. 2018a. Important juvenile manta ray habitat at Flower Garden Banks National Marine Sanctuary in the northwestern Gulf of Mexico. Marine Biology **165**(7):111.

Stewart JD, et al. 2018b. Research Priorities to Support Effective Manta and Devil Ray Conservation. Frontiers in Marine Science **5**:314.

Sukumaran J and Knowles LL. 2017. Multispecies coalescent delimits structure, not specie. Proceedings of the National Academy of Sciences **114**(7):1607–1612.

Walter RP, et al. 2014. First record of living Manta alfredi × Manta birostris hybrid. Marine Biodiversity **44**(1):1–2.

White WT, Corrigan S, Yang L, Henderson AC, Bazinet AL, Swofford DL and Naylor GJP. 2017. Phylogeny of the manta and devilrays (Chondrichthyes: Mobulidae), with an updated taxonomic arrangement for the family. Zoological Journal of the Linnean Society **182**(1):50-75.
